# Gaps and complex structurally variant loci in phased genome assemblies

**DOI:** 10.1101/2022.07.06.498874

**Authors:** David Porubsky, Mitchell R. Vollger, William T. Harvey, Allison N. Rozanski, Peter Ebert, Glenn Hickey, Patrick Hasenfeld, Ashley D. Sanders, Catherine Stober, The Human Pangenome Reference Consortium, Jan O. Korbel, Benedict Paten, Tobias Marschall, Evan E. Eichler

**Author notes:** List of contributing consortium members (The Human Pangenome Reference Consortium).

## Abstract

There has been tremendous progress in the production of phased genome assemblies by combining long-read data with parental information or linking read data. Nevertheless, a typical phased genome assembly generated by trio-hifiasm still generates more than ~140 gaps. We perform a detailed analysis of gaps, assembly breaks, and misorientations from 77 phased and assembled human genomes (154 unique haplotypes). We find that trio-based approaches using HiFi are the current gold standard although chromosome-wide phasing accuracy is comparable when using Strand-seq instead of parental data. We find two-thirds of defined contig ends cluster near the largest and most identical repeats [including segmental duplications (35.4%) or satellite DNA (22.3%) or to regions enriched in GA/AT rich DNA (27.4%)]. As a result, 1513 protein-coding genes overlap assembly gaps in at least one haplotype and 231 are recurrently disrupted or missing from five or more haplotypes. In addition, we estimate that 6-7 Mbp of DNA are incorrectly orientated per haplotype irrespective of whether trio-free or trio-based approaches are employed. 81% of such misorientations correspond to *bona fide* large inversion polymorphisms in the human species, most of which are flanked by large identical segmental duplications. In addition, we also identify large-scale alignment discontinuities consistent with an 11.9 Mbp deletion and 161.4 Mbp of insertion per human haploid genome. While 99% of this variation corresponds to satellite DNA, we identify 230 regions of the euchromatic DNA with frequent expansions and contractions, nearly half of which overlap with 197 protein-coding genes. Although not completely resolved, these regions include copy number polymorphic and biomedically relevant genic regions where complete resolution and a pangenome representation will be most useful, yet most challenging, to realize.

## INTRODUCTION

Assembly gaps are, unfortunately, still an integral feature of every *de novo* genome assembly. This *status quo* will remain until the sequencing technology and assembly algorithms evolve so that each homologous chromosome of any genome can be routinely assembled telomere-to-telomere (T2T) in an automated fashion. Key to this aspirational goal is understanding why gaps persist, which in turn requires a detailed analysis of gap size, frequency, genomic location, and the sequence properties that define these regions. The last two years have witnessed tremendous progress with respect to advances in sequencing technology (Lu, Giordano, and Ning 2016; Wenger et al. 2019; Vollger et al. 2019) as well as numerous assembly strategies that now make it possible to phase and assemble >95% of the content of a diploid genome (Logsdon et al. 2021; Jarvis et al. 2022). In particular, the development of HiFi (high-fidelity) Pacific Biosciences based on circular consensus sequencing (CCS) provides ~20 kbp sequencing reads that now rival short reads with respect to their accuracy (QV>40), while the Oxford Nanopore Technologies (ONT) platform now can generate sequencing reads in excess of 100 kbp (so called ultra-long (UL) sequencing reads) (Shafin et al. 2020; Nurk et al. 2020; Logsdon et al. 2021). The use of parent–child trio (trio-hifiasm) Illumina whole-genome sequencing (WGS) data provides the greatest power to phase a genome into its constituent paternal and maternal haplotypes. In the absence of parental data, however, methods have been developed (PGAS and HiC-hifiasm) using long-range linked data such as Strand-seq (Porubsky et al. 2021) or Hi-C (Garg et al. 2021; Kronenberg et al. 2021; Cheng et al. 2022) that can phase genomes locally as well as at the chromosomal level.

As a result of these developments, genome assemblies have changed in two significant ways over the last two years. We no longer consider collapsed 3 Gbp genome assemblies as state-of-the-art (i.e., one representation of an individual where both haplotypes are merged) but instead consider two genomes for every diploid genome assembled (i.e., 6 Gbp vs. 3 Gbp) where parental haplotypes are phased and fully resolved. Secondly and, in part because of the first, the number of gaps being produced has reduced from thousands to only a few hundred. With the completion and annotation of the first T2T genome (Nurk et al. 2022), we are in a position to characterize the properties of the gaps that remain when diploid human genomes are routinely sequenced. We focus on a detailed characterization of these remaining gaps in an effort to understand their origin, biology, and the relative importance of getting these through the last impasses to T2T assembly. We focus on human diploid genomes, since resolution of the gaps will improve discovery of both disease-related variation as well as genetic changes important for the evolution and adaptation of our species.

## RESULTS

We investigated the gaps and contig breaks in a total of 182 haploid assemblies obtained from a diversity panel of 77 unique human samples sequenced with long-read technology. The underlying long-read data and assemblies were generated by two consortia over the last two years, the Human Genome Structural Variation Consortium (HGSVC, 88 assemblies) and Human Pangenome Reference Consortium (HPRC, 94 assemblies), using different long-read sequencing platforms as well as assembly strategies. The HGSVC employed two different long-read sequencing technologies, continuous long read (CLR, 60 assemblies) (Ebert et al. 2021) and CCS (or high-fidelity ‘HiFi’ sequencing, 28 assemblies) with an additional eight samples shared between HGSCV and HPRC used only for validation purposes. CCS and CLR data from HGSVC were assembled using a trio-free assembly pipeline, called PGAS (Porubsky et al. 2021; Ebert et al. 2021; Ebler et al. 2022) employing both the Peregrine (Chin and Khalak 2019) (PGASv12) and the hifiasm (Cheng et al. 2021) (PGASv13) assembler for CCS and the Flye assembler (Kolmogorov et al. 2019) for CLR reads. The HPRC effort, which began more than a year later, focused exclusively on CCS data (n=94) generated from diploid samples assembled using trio-based hifiasm (Cheng et al. 2021). Here, parent–child data were directly used to aid assembly phasing of all HPRC samples (Wang et al. 2022; Liao et al. 2022) allowing for both platform and methodology comparisons.

### Evaluation metrics and gap definitions

In this study, we set out to evaluate assembly quality and completeness using four metrics (**Methods**). We start with defining regions between subsequent contigs mapped to the T2T human genome reference. These are defined based on reliable ‘contig end alignments’ (≥50 kbp at the contig edges) mapped in agreement with an expected contig length. Contig end alignments were used to localize regions (assembly gaps) in between subsequent contigs (**Fig. 1A, i**). Second, we define ‘simple contig ends’ as terminal contig positions with respect to the reference genome. Simple contig ends were used for enrichment analysis of various genomic features nearby terminal contig alignment positions (**Fig. 1A, ii**). To evaluate structural differences between assemblies we set to document all regions that break contig alignments, referred to here as ‘contig alignments discontinuities’. We focus on discontinuities that create internal gaps within contig alignments less than 1 Mbp in length to document regions of putative structural differences that cannot be readily aligned to a single reference (**Fig. 1A, iii**). Lastly, we turn our attention to regions with a higher coverage than expected in a haploid genome (multi-coverage regions) caused by two or more overlapping contig alignments. Such regions point to positions of either true structural differences or genome assembly artifacts (**Fig. 1A, iv**).

**Figure 1:**
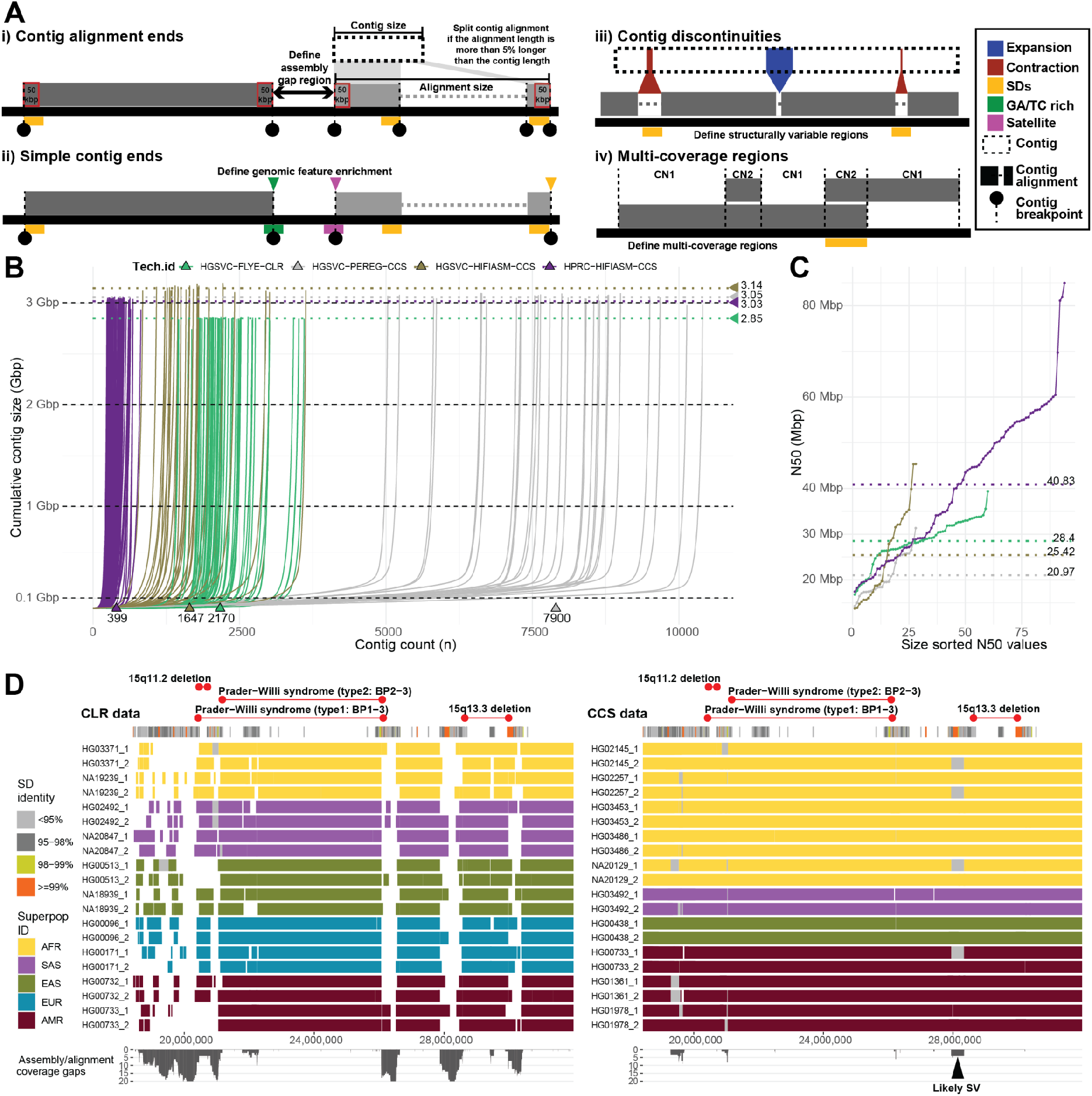
Comparison and evaluation of phased assemblies. **A**) Assembly metrics evaluated in this study: **i**) Contig alignment ends are defined as terminal contig alignments such that the total alignment size does not exceed the actual contig size by more than 5%. When this requirement is not met, multiple contig end alignments will be reported. **ii**) Simple contig ends are defined as the first and last alignments of each contig to the reference (T2T-CHM13 v1.1) of at least 10-50 kbp in size. **iii**) Contig discontinuities are defined as alignment gaps between subsequent pieces of a single contig smaller than 1 Mbp. **iv**) Detection of regions with coverage more than 1 as is expected for a haploid genome. **B**) A cumulative contig size distribution colored by assembly technology. Each line represents a single haploid assembly (HGSVC-FLYE-CLR (n=60), HGSVC-PEREG-CCS (n=28), HGSVC-HIFIASM-CCS (n=28), HPRC-HIFIASM-CCS (n=94)). Median total assembly length per assembly technology is highlighted as horizontal dotted lines. **C**) Contig N50 values colored by assembly technology in B). Each dot represents a single haploid assembly. Median N50 value per assembly technology is highlighted as horizontal dotted lines. **D**) Track definition from top to bottom: Regions corresponding to known genomic disorders between 15q11.2-15q13.3. Below is the annotation of SDs in this region colored by sequence identity. Main track shows the visualization of contig alignments for 10 random samples from trio-free CLR assemblies (left) in comparison to trio-based HPRC assemblies (right). Contig alignments are colored by sample superpopulation. White spaces between contig alignments represent boundaries between subsequent contig. Spaces filled with gray color represent unaligned portions of a single contig in respect to the reference (T2T-CHM13) and likely represents a structural variation (black arrowhead). Last track summarizes the extent of assembly (between contigm, white space) gaps and within contig gaps (gray rectangles) as coverage plot.

### Platform and assembly method comparisons

We initially compared assembly statistics between different sequencing technologies and assembly algorithms to determine what combination provides the most continuous and complete assembly. The most fragmented assemblies were obtained using a combination of the trio-free PGAS pipeline and the Peregrine assembler with a median contig count of 7900 per assembly (Ebert et al. 2021). Improved contiguity was achieved by combination of the PGAS pipeline and CLR reads assembled by Flye (median contigs: 2170) and CCS reads assembled by hifiasm (median contigs: 1647) (Ebert et al. 2021); (Ebler et al. 2020). The most continuous assemblies were obtained using the trio-based hifiasm assembly resulting in an order of magnitude fewer gaps (e.g., 399 median contigs per assembly (**Fig. 1B**). The least complete assemblies resulted from a combination of PGAS and CLR data (median size 2.85 Gbp). This is expected as comparably higher error rates of CLR in comparison to CCS reads prevent them from assembling highly identical segmental duplications (SDs) in the human genome. Assemblies using CCS reads provide comparable assembly completeness (median size ~3.05 Gbp) with a slightly higher median assembly size for the trio-free PGAS pipeline combined with hifiasm (median size 3.14 Gbp) (**Fig. 1B**). Lastly, the assembly contiguity was evaluated as a function of contig N50, and again we conclude that trio-based assembly (N50: 40.83 Mbp) outperforms those assembled in trio-free settings (**Fig. 1C**). Due to suboptimal performance, we excluded Peregrine assemblies from subsequent analyses.

Consistent with recent studies (Jarvis et al. 2022), trio-based assemblies contain the least number of gaps between contig alignment ends (median: 141) followed by PGAS-hifiasm with about double that amount (median: 320) and PGAS-Flye (median: 392) (**Fig. S1A**). Based on projections to the T2T-CHM13, the number of missing base pairs follows a similar trend with trio-based assemblies having the least number of bases within gaps between defined contig alignment ends (median: 78.4 Mbp) followed by PGAS-hifiasm (median: 126.7 Mbp) and PGAS-Flye (median: 244.8 Mbp) (**Fig. S1B**). There are two obvious outliers in trio-based assemblies with very long gaps caused by contigs mapping partly to PAR1 region and a q-arm of chromosome X in two paternal haplotypes (HG02055, HG02572) (**Fig. S2**).

Superior quality of trio-based assemblies (HPRC) is partly ensured by better data quality with longer insert sizes and overall coverage in respect to data used in PGAS assemblies (HGSVC). Despite the longest insert sizes and the highest coverage, we observe that Flye assemblies of CLR have some of the greatest difficulty assembling complex SD regions due the comparably lower sequence accuracy (Wenger et al. 2019) (**Fig. S1C**). This shortcoming is especially problematic in disease-relevant regions such as the Prader-Willi critical region (15q11.2-15q13.3) where highly identical SDs are largely absent from CLR-based assemblies (**Fig. 1D, white gaps**). This results in about 59.9 Mbp missing base pairs in the presented CLR assemblies, which is in stark contrast to only ~690 kbp missing base pairs in the CCS-based assemblies. In addition, this prevents us from correctly evaluating recently reported heterozygous inversion in this region in sample HG02492 (Porubsky et al. 2022). As a result, the more contiguous CCS-based assemblies allow us to begin to assess copy number and structural variation with respect to the reference genome (**Fig. 1D, gray gaps**). Given these observations, we exclude CLR-based assemblies from subsequent analysis and focus exclusively on CCS-hifiasm assemblies.

### Parent child trio-based vs. trio-free assemblies

We compared in more detail eight human genomes where both long-range linked reads (Strand-seq) and parental data (Illumina WGS) were available from the same individuals. Using the same underlying long-read input data (CCS), we specifically performed a head-to-head comparison of trio-based (TRIO) and trio-free (PGAS) assemblies. We find that assemblies generated in the absence of parental data (trio-free) have about twice as many contigs and a decreased contig N50 by ~10 Mbp (**Fig. S3**). We next evaluated phasing accuracy of trio-free assemblies using the genomes phased by parental data as the truth set (**Methods**). For the metacentric and submetacentric chromosomes we observe high accuracy of phased 1 Mbp segments achieving 98% concordance with trio-based phasing. With acrocentric chromosomes this accuracy drops to 94% (**Fig. 2A, Fig. S4**). The majority of incorrectly assigned 1 Mbp segments (>75%) map within centromeric satellite repeats (**Fig. S5**). There was only one sample (HG01891) with large-scale switch errors on a short arm of chromosome 9 (~42 Mbp) and one at the very end of chromosome 9 (~1 Mbp) (**Fig. 2B**). The data shows that trio-free assemblies provide comparable phasing accuracy and completeness and are a viable option for phased genome assembly for samples where parental data are not easily available or are cost prohibitive.

**Figure 2:**
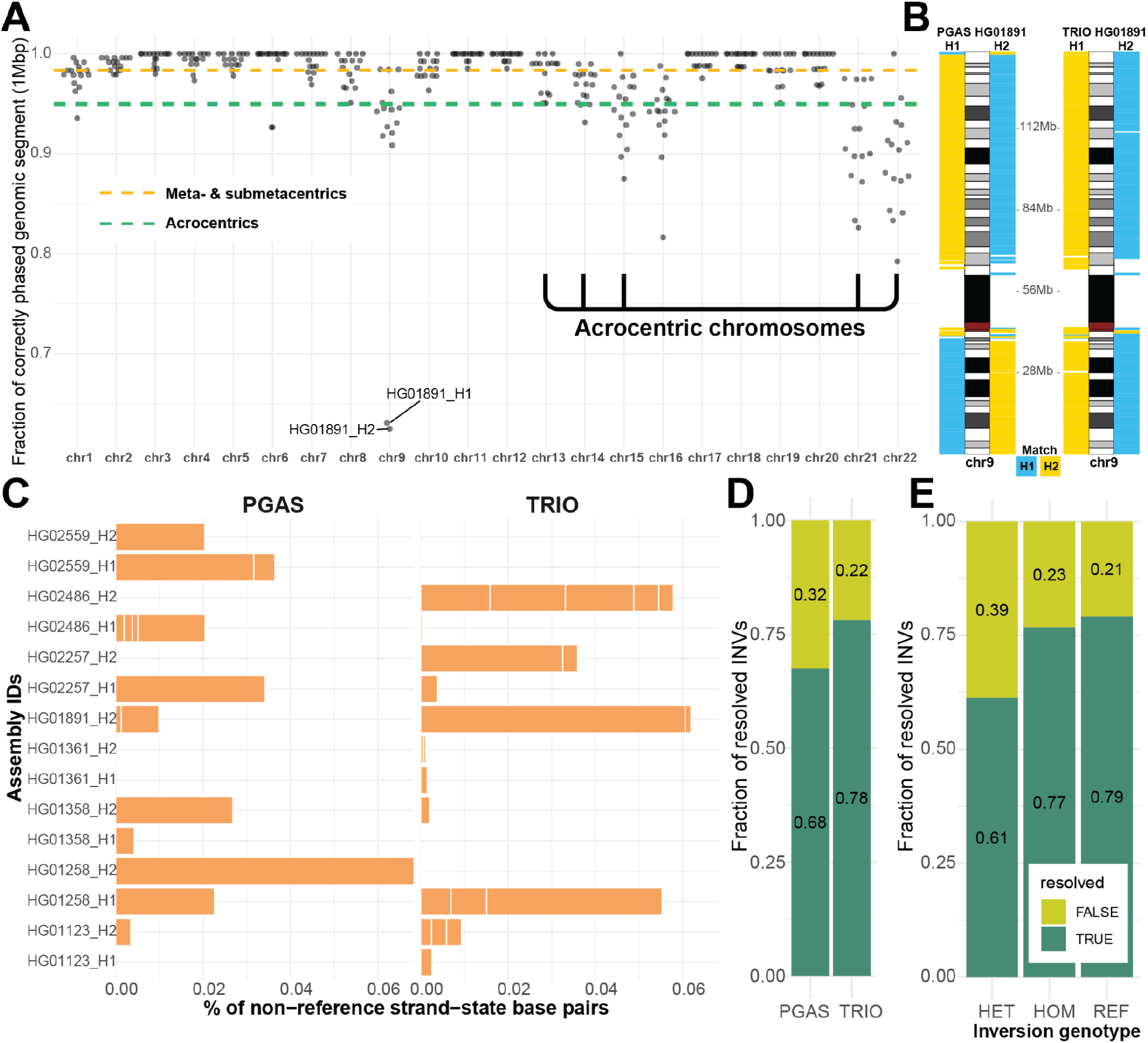
Phasing accuracy and inversion analysis of trio-based and trio-free assemblies. **A**) Phasing accuracy of PGAS (trio-free) assemblies in respect to trio-based phasing. **B**) Haplotype assignment of 1 Mbp-sized blocks (left from ideogram - H1, right from ideogram - H2) to either haplotype 1 or 2 (H1 - blue, H2 - yellow) using single-nucleotide polymorphisms phased using trio information (1KG panel) with respect to the reference (GRCh38). **C**) A barplot reporting the percentage of base pairs in an opposite orientation than expected based on Strand-seq analysis of assembly directionality, shown separately for trio-free (PGAS, left, n=15) and trio-based (TRIO, right, n=23) assemblies. **D**) Fraction of tested inversion sites that are fully informative (TRUE, dark green). **E**) Fraction of tested inversion sites that are fully informative (TRUE, dark green) as a function of inversion genotype (HET - heterozygous, HOM - homozygous inverted, REF - homozygous reference).

Strand-seq also preserves directionality of single-stranded DNA and thus is also able to unambiguously define misoriented regions of the genome. Such misorientations will appear as unresolved homozygous inversions based on Strand-seq reads mapping from the original genome sample (**Methods**). Surprisingly, we detected comparable numbers of unresolved homozygous inverted regions in trio-based (n=23) and trio-free assemblies (n=15), respectively (**Table S1**) resulting in 6.8 Mbp (0.23%) and 7.3 Mbp (0.25%) of misoriented base pairs per assembly (**Fig. 2C**). The majority (31/38, >81%) of these misorientations overlap with previously defined true inversion polymorphisms in the human genome (Porubsky et al. 2022), six of which were unresolved in both trio-based and trio-free assemblies (**Fig. S6A**) and, as expected, are flanked by large tracts of SDs (**Fig. S6B**). Some of these span genomic disorder critical regions where recurrent *de novo* CNVs associate with neurodevelopmental delay, such as the 16p11.2-p12.2 microdeletion and microduplication syndrome (**Fig. S7**).

We more systematically evaluated the potential of both assembly approaches to resolve known large (≥100 kbp, n=20) inversions considering both heterozygous as well as homozygous sites (**Methods**). Trio-based assemblies resolve 78% of inversion polymorphisms while trio-free assemblies resolve 68% (**Fig. 2D**). Trio-based approaches generally more accurately represent more inverted base pairs (64%) when compared to the trio-free approach (48%) (**Fig. S8B**) by virtue of the fact they often assemble one end of an inversion polymorphism (**Fig. S8A**). It is noteworthy that nearly a quarter of all large inversion polymorphisms are not accurately represented in existing trio-based genome assemblies with heterozygous inversions being the most difficult to fully resolve (**Fig. 2E, Fig. S8C**). All sites (n=14) that are unresolved two and more times in trio-based and trio-free assemblies are flanked by large (>40 kbp, median: 228.2 kbp) highly identical SDs (median 99.4%). The availability of Strand-seq data provides a valuable orthogonal method for detection of such errors in the assembly which in turn can guide targeted reassemblies of such regions using ultra-long ONT reads.

### Sequence properties of the gaps

Because the HPRC-phased genome assemblies represent the current state-of-the-art in terms of both accuracy, phasing and contiguity (**Fig. 1 and Fig. 2**), we focused on a more in-depth analysis of sequence content of gap regions by mapping all sequence contigs to the complete human reference (T2T-CHM13, v1.1) (Nurk et al. 2022). Among the 94 HPRC haplotype assemblies, we identified a total of 68,515 simple contig ends for an average of 729 per haplotype (median of 700) (**Fig. 1C, Table S2**). Of these contig breaks, about two-thirds correspond to SDs [35.4% (11702+12550)/68515] or satellite DNA [22.3%, (2896+12363)/68515] (**Fig. 3A**). Because long tracts of GA repeats have been predicted to reduce the coverage of CCS data (Nurk et al. 2020), it is important to note that 27.4% [(6212+12550)/68515] of the gaps, including recurring gaps, within the assemblies correspond to regions where high GA/TC tracts are observed (1 kbp window with more than 80% GA/TC within 10 kbp). These GA/TC tracts show the most substantial (29.36-fold) (**Fig. 3B**) enrichment for gaps, and along with high AT content account for ~40% of the assembly breaks not associated with large repetitive sequences [(6212+5494)/(68515-2896-12363-11702-12550)]. Controlling for sequence coverage, we estimate that nearly two-thirds of the GA/TC gaps can be remedied by simply increasing sequence coverage from ~30-fold to 50-fold (**Fig. 3C**). However, we also find long tracts of GA/TC repeats also associate non-randomly with regions of SDs (**Fig. 3A**). In such regions, increasing coverage has little effect on reducing the number of gaps and perhaps even the opposite effect (**Fig. 3D**). We considered both the length and sequence identity of SDs and found that the longer and more identical a SD is, the more likely it was associated with a gap (**Fig. 3E, Fig. S9**). Thus, the longest and most identical SDs are preferentially associated with gaps in the majority of analyzed assemblies (**Fig. 3E, Fig. S9**).

**Figure 3:**
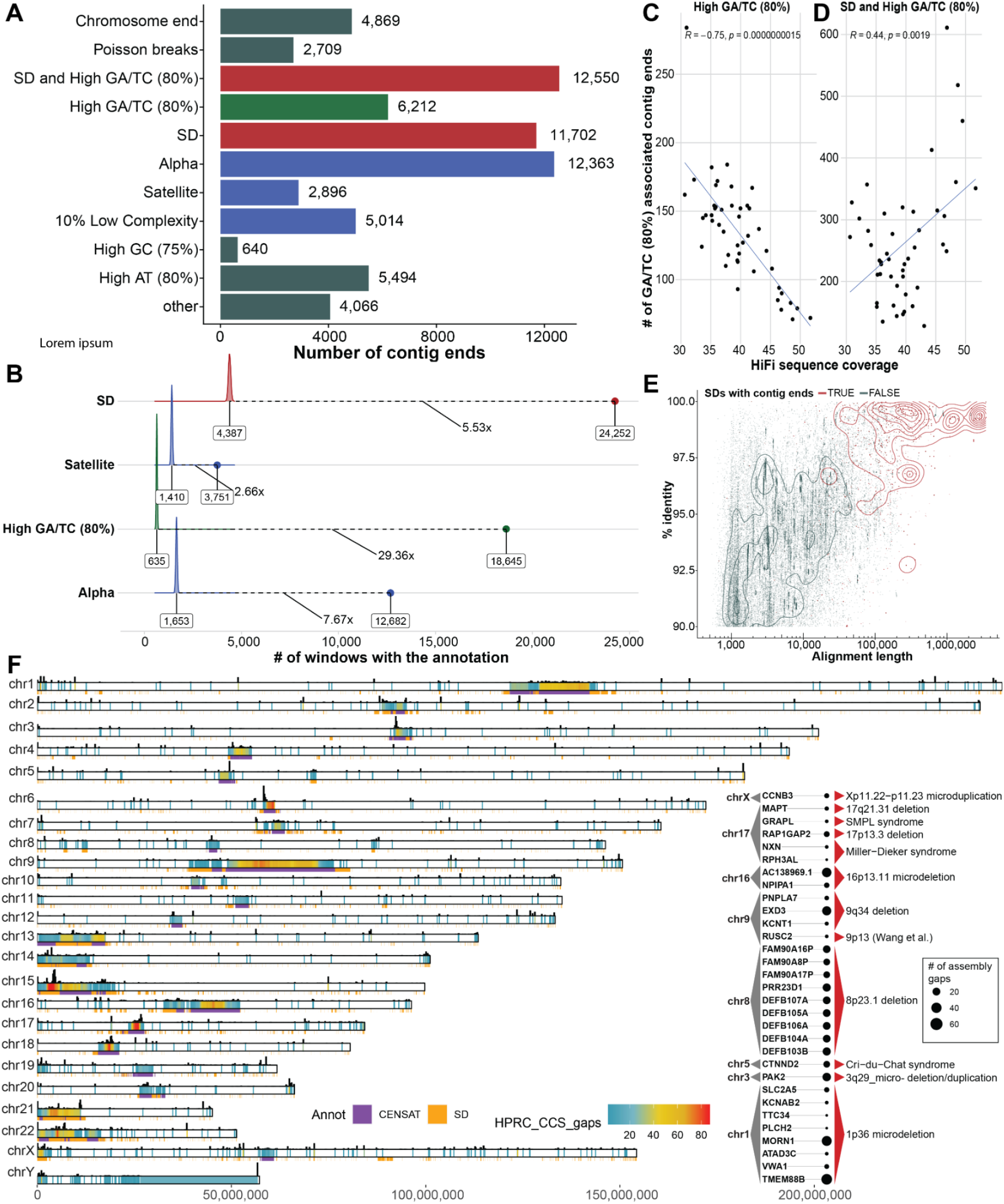
Sequence properties at defined contig ends. **A**) The number of simple contig ends that are within or near (at most 10 kbp) a particular sequence annotation. Annotations are nonredundant and are prioritized in the order shown, e.g., if a contig end is near the end of a chromosome and in SD it will only be annotated as a chromosome end. NOTE: Chromosome ends are contig ends within the last 100 kbp of contigs. Poisson ends are contig ends that happen in only 1 haplotype (nonrecurrent and therefore likely to be random). SD and High GA/TC mean that the end is within 10 kbp of an SD and within 10 kbp of a 1 kbp window with at least 80% GA/TC content. **B**) Fold enrichment in the number of contigs ends within 10 kbp of a sequence annotation compared to a distribution of randomly placed contig end simulations (10,000 permutations). Shown in text is the median of the random distribution (left), the fold enrichment (middle), and the observed value (right). In this analysis contig ends may exist in multiple categories, e.g. if a contig end is near both an SD and a satellite sequence it will appear in both simulations. **C**) Effect of HiFi coverage on number of GA/TC breaks. Negatively correlated when considered independently; however, when combined with SD, the trend is inverted as shown in **D. E**) All SDs in T2T-CHM13 displayed by their length and % identity (blue) vs the SDs that intersect contig ends (red). 02492. As **F**) Genome-wide distribution of gaps defined in between contig alignment ends (**Methods**) across all HPRC assemblies (n=94). Color range reflects the number of assembly gaps overlapping each other in any given genomic region. On the top of each chromosomal bar there is a density of simple contig ends. The height of each bar reflects a number of simple contig ends counted in 200 kbp long genomic bins.

Despite the differences in contig end definition, we found a high level of agreement between simple contig ends and assembly gap regions with >85% of simple contig ends falling into assembly gaps and >99% of assembly gaps overlapping with simple contig ends (**Fig. 3F, Fig. S10**). Assembly gaps are regions that are not completely assembled across HPRC assemblies. This is especially problematic when assessing human diversity among protein-coding genes. The whole set of assembly gaps (n=14662) from all HPRC assemblies overlaps a total of 1513 protein-coding genes (**Fig. S11**) that falls within 894 nonredundant gap regions. There are 231 protein-coding genes that fall within regions broken in five or more HPRC assemblies (**Fig. S12, Table S3**) and 31 of these lie within regions of recurrent microdeletion and microduplication syndromes (Cooper et al. 2011; Coe et al. 2014). Among these, there are a number of biomedically relevant genes such as *PAK2* affected by 3q29 microdeletion, *CTNND2* affected in Cri-du-Chat syndrome, or *MAPT* affected by 17q21.31 microdeletion (**Fig. 3F, inset**).

Overall, we define 592 nonredundant regions, outside of satellite DNA, with an assembly gap in five or more of the HPRC assemblies (**Fig. S13, Table S4**). Among the most recurrent gaps, there are 44 euchromatic regions which fail to resolve in half or more of the HPRC assemblies. While a third of these are associated with SDs, 28 of these are dropouts associated with the presence of low-complexity DNA (**Table S5**). In these regions we observe continuous tracts of dinucleotides (AT or GA/TC) ranging from ~300-6500 kbp (**Fig. S14**); however, we noticed a number of such low-complexity tracts in regions associated with SDs (n=16) (**Fig. S15**). We further explored the extent of variability in size of these low-complexity regions between humans and nonhuman primates. We catalog 27/44 regions with observable differences in size of dinucleotide tracts, with humans carrying longer dinucleotide tracts in all but one instance (**Fig. 4A**). Our analysis suggests that many of these regions appear to have expanded specifically in the human lineage where they continue to show variability in size (**Fig. 4B,C**).

**Figure 4:**
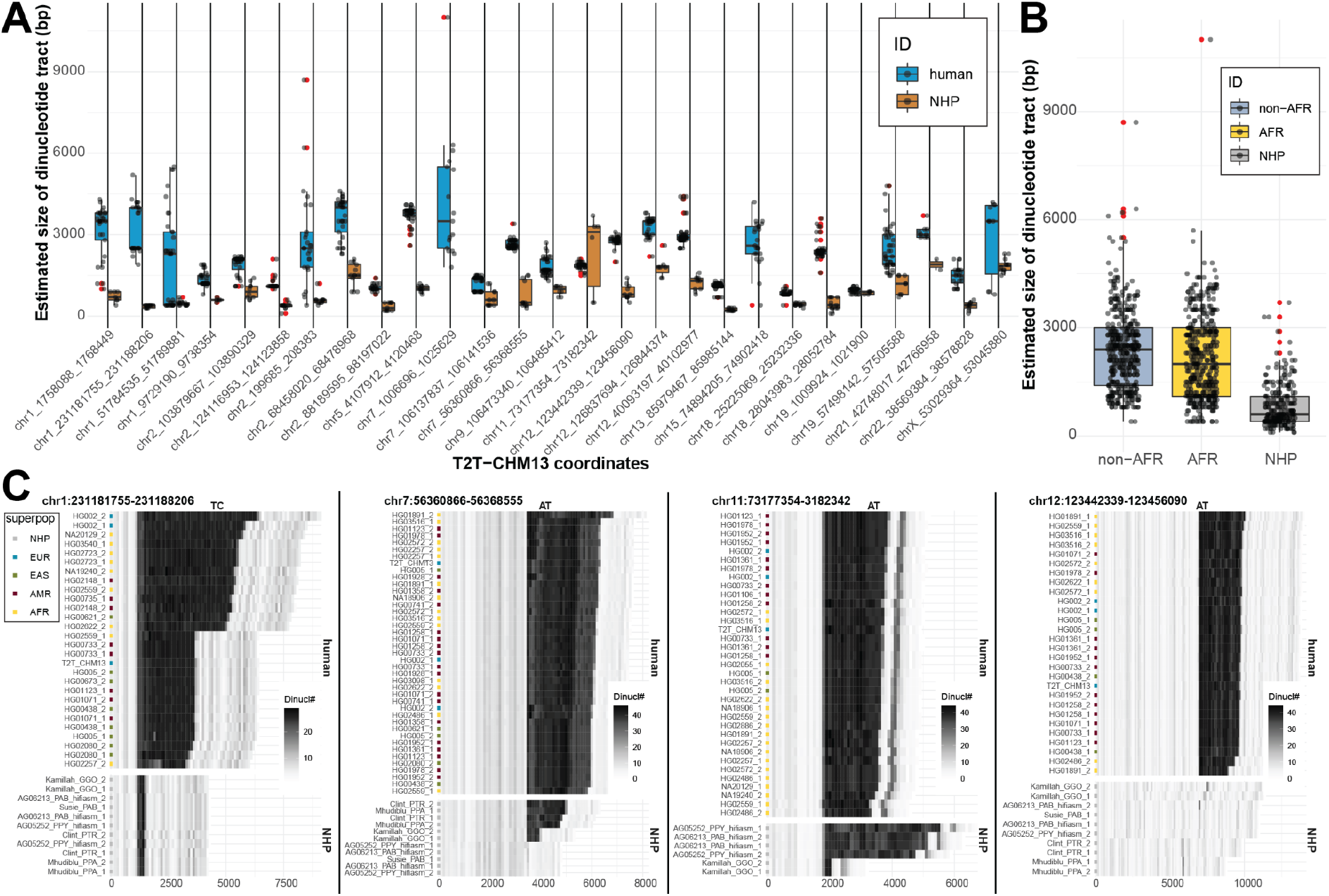
Sequence variation in low-complexity regions. **A**) Comparison of size distribution of dinucleotide tracts (y-axis) between human (blue) and nonhuman primates (NHP, brown) for 27 selected regions (**Methods**). Outliers are highlighted as red dots. **B**) A summary of size distribution of dinucleotide tracts (y-axis) between human samples of African (AFR, yellow) and non-African (non-AFR, light blue) origin, and nonhuman primates (NHP, gray) across all complete assemblies from 27 selected regions. **C**) Difference in dinucleotide frequency (TC, AT) between humans and nonhuman primates (NHP) in four genomic regions. Shades of gray color reflect the number of detected dinucleotides (defined at the top of each plot) in 100bp long DNA sequence chunks. Assembly names (y-axis) from NHPs contain sample IDs and species specific ID (PTR - Pan Troglodytes, GGO - Gorilla Gorilla, PPA - Pan Paniscus, MMU - Macaca Mulatta PAB - Pongo Abelii, PPY - Pongo Pygmaeus). Numbers 1 and 2 represent parental homolog IDs of given sample assembly.

### Discontinuous alignments and large structural variants

One of the advantages of the new assemblies of the human genome is that they are not guided by existing human references. Such *de novo* assemblies have the potential to identify large discontinuities corresponding to potential larger forms of genetic variation including partially sequence-resolved copy number variants (CNVs). We searched specifically for contig alignment discontinuities (<1 Mbp) as identified by alignment to the complete human reference genome (T2T-CHM13, v1.1) (**Fig. 1A, Methods**). Across all 94 human haplotypes we report a median 6.6% and 0.06% of unaligned bases per assembly within and outside of centromeric satellite DNA, respectively (**Fig. S16**). Per haploid genome we define a median number of 165 contractions and 262 expansions, which corresponds to about 11.9 Mbp and 161.4 Mbp, respectively (**Fig. S17A-B**). The vast majority of these bases (contractions 10.9 Mbp, expansions 159.8 Mbp) belong to centromeric satellite DNA which is known to vary extensively in size and composition among human haplotypes and is often incompletely assembled (**Fig. S17C**). Nevertheless, within euchromatic regions we identified 230 regions that showed evidence of contraction (n=120) or expansion (n=110) in multiple human haplotypes (≥5) when compared to the T2T-CHM13 reference (**Fig. 5A, Table S6**). A large number of these regions overlap with SDs ~40% (93/230) and include biomedically relevant loci that are known to be structurally variable such as 8p23.1, HLA, *SMN1/SMN2* and *TBC1D3* (Vollger, Guitart, et al. 2022) (**Fig. 5B, Fig. S18**). Based on the read-depth analysis of Illumina WGS data we confirm 41 of these regions, the majority of which correspond to copy number losses in their respective genomes (**Fig. S19, Methods**). We highlight a region on chromosome 11 (chr11:55535304-55628574, 11q12.1) where the contracted region (~93 kbp) is associated with short inversion (~4 kbp) that flips *OR4C6* into a direct orientation in respect to *OR4C11*, which likely promotes a microdeletion via NAHR as this deletion is observed in association with inverted haplotype (**Fig. S20**).

**Figure 5:**
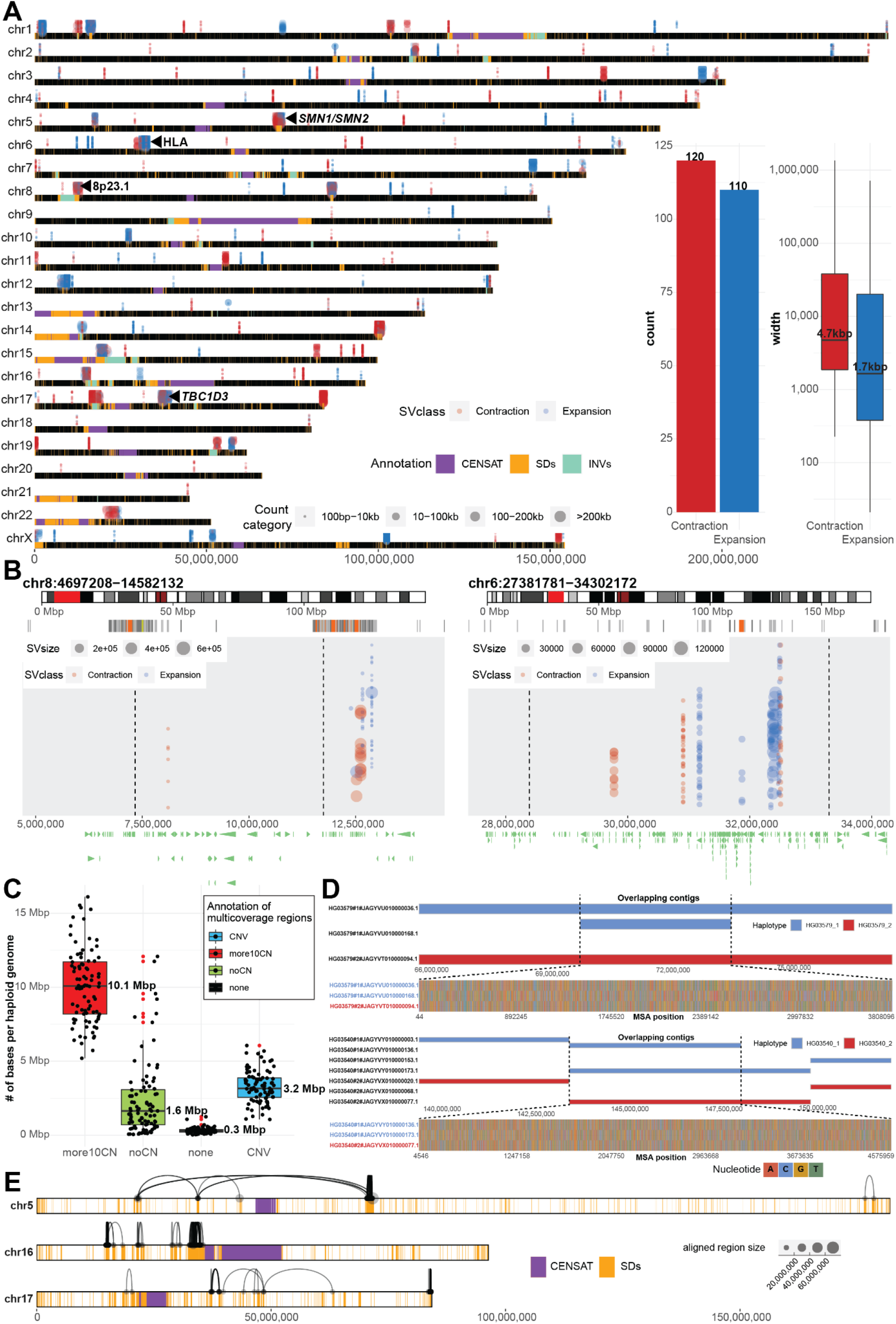
Tracking contig alignment discontinuities and multi-coverage regions. **A**) Genome-wide distribution of frequent (n=230) contig alignments discontinuities (1 kbp - 1 Mbp in size). Each gap is represented in each separate assembly (HPRC - 94, HGSVC - 28) by a colored dot (blue - INS - expansion, red - DEL - contraction) and the size of each dot represents the size of the event in contig coordinates. A region is defined as an expansion (INS - blue) if there is a gap in a contig alignment (in reference T2T-CHM13, v1.1 coordinates) that is smaller than the sequence within a contig itself delineated by the left and right alignment flanking the gap. In contrast a contraction (DEL - red) is defined as a gap in a contig alignment (in reference T2T-CHM13, v1.1 coordinates) that is larger than the sequence within a contig itself delineated by the left and right alignment around the gap. Putative expansions and contraction above the horizontal chromosomal lines were detected in HPRC assemblies and those below the lines in HGSVC assemblies. Centromeric satellite regions are highlighted by gray rectangles and regions of segmental duplications (SDs) as orange rectangles on top of each chromosomal line (black). **B**) Example regions (Left: Defensin locus - 8p23.1; right - HLA locus) with frequent expansions and contractions. Each region is highlighted as a red rectangle on chromosome-specific ideogram (top track). Below there is an SD annotation for a given region represented as a set of rectangles colored by sequence identity. Expansions and contractions of each contig alignment in respect to the reference (T2T-CHM13, v1.1) are depicted as blue and red dots, respectively. Size of each dot represents the size of an event. **C**) Assignment of total number base pairs covered by multiple contig alignments, in each haploid genome (n=88), into four categories based on agreement with short-read-based CNV profiles. **D**) An example regions in sample HG03579 and HG03540 where there are overlapping contigs associated with loss of heterozygosity. Top track shows contig alignments in a given region separately for haplotype 1 (blue, paternal) and haplotype 2 (red, maternal). Overlapping contig alignments are stacked on top of each other. Bottom track shows all variable positions detected in multiple sequence alignment (MSA) over the region where contigs overlap (dashed lines). Here one of the paternal contig is nearly identical to maternal contig over the contig overlap. **E**) Chromosomes 5, 16 and 17 are depicted as horizontal bars with the locations of SDs and centromeric regions highlighted as orange and purple rectangles, respectively. Contig alignment ends divided into multiple pieces are visualized as links between subsequent pieces of a single contig aligned to the reference (T2T-CHM13 v1.1). Length of the aligned pieces of a contig are defined by the size of each dot.

In addition to the assembled sequence that does not readily map to the reference, we also cataloged regions where there are multiple contig mappings (>1) instead of the expected haploid single copy (**Fig. 1A, iv**). Per haplotype we observe ~15.4 Mbp of euchromatic sequence with multiple contig mappings in respect to the reference (T2T-CHM13, v1.1). While such multimapping regions likely represent CNV regions arising from SD, they may also result from ambiguous contig mappings or artifacts of the assembly process. To enrich for true CNVs, we searched for CNV regions that were also supported by read-depth analysis of short-read data (**Methods**). Indeed, we identified ~3.2 Mbp predicted to be CNV (2-10 copies) and being supported by short read sequence data. An even greater fraction (~10.1 Mbp) of multimapping regions show greater CNVs (>10 copies) based on short-read depth although the true copy number is more difficult to determine as majority (>95%) of these regions overlap with SDs by more than 90%

Nevertheless, we identify ~1.6 Mbp per haplotype of multimapping regions where we find no obvious CNV in short-read data (**Fig. 5C**). We note that a subset of these are large (≥500 kbp) and often (85/118) represent sequence contigs that are completely embedded within another larger contig in a single haplotype. We investigated eight of the longest such contigs in more detail (**Methods**). Comparison of heterozygous single-nucleotide variant (SNV) patterns across these regions based on CCS reads (deepvariant calls) and phased assemblies (dipcalls) reveals conspicuous stretches of loss of heterozygosity over the region where the multimapping contigs overlap (**Fig. S21**). Closer inspection reveals that the sequence variation between parental haplotypes is, however, not lost but rather is present only in one contig while the other contig is nearly identical to the other parental haplotype (**Fig. 5D**). While the origin of such assembly artifacts is unclear, such overlapping contigs will likely pose challenges for SNV calling depending on which, if any, sequence contig is chosen.

We focussed specifically on euchromatic regions where both long-read and short-read data were in agreement regarding increased CNV (<10 copy number increase with respect to the reference). We identified 255 nonredundant CNV regions that encompass 44.9 Mbp of the genome (**Table S7**). Of these CNV regions, 87% (39.1 Mbp) correspond to SDs that are known to be copy number variable because of their propensity to undergo NAHR (Sharp et al. 2006; Sudmant et al. 2010, 2015) (**Fig. S22**). We find that genomes of African ancestry carry more CNV bases (~3.5 Mbp) when compared to other non-African populations (**Fig. S23**) consistent with previous reports (Sudmant et al. 2015; Chaisson et al. 2019; Byrska-Bishop et al., n.d.). The regions are particularly gene-rich and we identify 420 protein-coding genes among 165 of them (**Table S7**).

Large-scale CNVs within an assembled contig may also lead to alignment discontinuities where contig alignment ends map far away from each other thus exceeding the expected contig length. We identified 1721 contigs whose alignments have exceeded the absolute contig length by more than 5% (**Fig. 1A, i**). While the majority of such contigs were observed in satellite DNA, we identified 391 contigs mapping outside of centromeric satellites of which ~98% are associated with SDs (**Fig. 5E, Fig. S24**). While we cannot exclude the possibility that such unusual patterns of homology result from assembly error or inability of mapping algorithms (such as minimap2) to distinguish between paralogous sequences due to high sequence identity (e.g., *SMN1/2* region (**Fig. S25**)), complete haplotype sequence and assembly of these regions is likely to provide new insights into patterns of human genetic variation and the mutational processes that shape them (Vollger, DeWitt, et al. 2022).

## DISCUSSION

The recently released gapless assembly of the first haploid human genome has set the bar for T2T human genome assemblies (Nurk et al. 2022). Extending this to diploid samples requires a detailed analysis of remaining gaps to guide new developments in both sequencing technology and assembly algorithms. Using multiple metrics, we provide a genome-wide assessment to characterize the nature of these last gaps of the human genome. There are several important conclusions. First, we show that recent improvements in sequencing technology (CLR versus CCS) and assembly algorithms (Peregrine versus hifiasm) reduce the number of gaps by ~3-fold. Second, the use of parental Illumina WGS data leads to gold standard phased genome assembly but the use of long-range linked-reads data such as Strand-seq can create phased assemblies with comparably low levels of switch error. Nevertheless, both trio-based and trio-free assemblies fail to correctly resolve the orientation of 6-7 Mbp of DNA. This is especially the case for large inversion polymorphisms that are flanked by high-identity SDs, which are one of the most difficult SV classes to accurately assemble (Chaisson et al. 2019; Porubsky et al. 2022). Such complex regions of the genome often coincide with morbid CNVs where the critical region toggles from a direct to an inverted configuration as a result of recurrent NAHR events (Porubsky et al. 2022).

The current state-of-the-art human diploid genome assembly is represented by ~140 gaps per haploid genome with about double the number when trio-free approaches, such as PGAS (Porubsky et al. 2021) are applied. Predictably, gaps cluster within copy number variable repeat-rich locations corresponding to the largest and most identical repeats (including satellites and SDs) or within low-complexity regions enriched in GA/AT dinucleotides. The latter results from a reduction in sequence coverage associated with the HiFi (CCS reads) sequencing platform over these particular motifs (Nurk et al. 2020). Interestingly, the degree of dropout shows some dependence on the size of the dinucleotide tracts with problematic regions ranging from 300 to 6.5 kbp in length. Many of these regions appear to have expanded specifically in the human–primate lineage so different regions are anticipated in other nonhuman genomes. Our analysis shows that increasing sequence coverage from 25-fold to 50-fold eliminates approximately two-thirds of such gaps. In contrast, increasing sequence coverage seems to have little effect in gaps associated with CNV SDs (**Fig. 3**). This is likely a consequence of the fact that insert size and sequence coverage are inversely correlated and, as a result, high-coverage samples suffer from smaller inserts that fail to resolve large SDs. In this regard, it is interesting that alternate long-read sequencing platforms such as ONT do not show the same inherent coverage biases toward GA/AT low-complexity repeats (Nurk et al. 2022). Coupled with their much longer read lengths (>50 kbp), we estimate that ~64% of the remaining gaps within HiFi assemblies can be traversed by ONT (**Fig. S26, Methods**). Approaches and assembly algorithms that couple both ONT and HiFi data (e.g., Verkko (Rautiainen et al. 2022)) may be necessary to close the remaining gaps and to achieve routine T2T assemblies of human genomes. The costs of generating deep long-read sequence coverage from two platforms to achieve a T2T human genome remain a significant cost (>$10,000) and throughput consideration.

One of the largest gains from T2T assemblies will be an improved understanding of human structural genetic diversity. While still incomplete, our analysis identifies ~6.6% and 0.06% of unaligned bases per haploid assembly localized within and outside of centromeric satellite DNA, respectively. Among such gaps caused by contig alignment discontinuities, we identify 230 regions that occurred in at least five haploid assemblies. Nearly half of these (~40%) map to SDs where variation and incomplete assembly pose particular challenges to alignment as well as interpretation. For example, within euchromatic regions, we identified ~15.4 Mbp of sequence per haplotype with two or more mappings per haplotype. Based on Illumina read-depth analysis, we estimate that 86% of these additional alignments represent *bona fide* human copy number variation. Nevertheless, ~1.6 Mbp of the reported extra alignments are likely false as there is no support in short-read data. Interestingly, such alignments are often represented by contigs embedded within other larger contigs where the overlapping contig alignments has lost allelic variation and now carry, instead, the allelic pattern of variation of the opposing parental haplotype. SNVs are, thus, still present but map to only one of the contigs generated by trio-hifiasm for a given haplotype. This is important because current variant-calling algorithms such as dipcall or pav tend to pick the longer more contiguous contig in both haploid assemblies to infer allelic variation. We predict that such artifacts may overestimate loss of heterozygosity regions when the longer contig devoid of SNVs is preferentially used.

A major challenge going forward will not only be to fully sequence resolve these regions but to represent complex SVs in such a way that they can be reliably interpreted and assayed in human genetic studies. One of the main objectives of the HPRC efforts is to project all human genome variation through a graph-based representation where every human haplotype represents a path in the graph. Unfortunately, there are regions in current genome assemblies that are still completely missing, incorrectly assembled, or otherwise pose challenges for the construction of such pangenome graphs. A set of regions, termed “brnn” regions, were identified and “trimmed” during the construction of the minigraph-cactus graph (Liao et al. 2022). These regions were excluded at least once but, in some instances, up to 88 times and mapped predictably to satellite DNA (n=149 regions or ~149.7 Mbp; ~28.9 Mbp in acrocentrics) and SD regions (n=301 regions or ~65.7 Mbp) but also correspond to protein-coding genes (n=171) as well as common inversion polymorphisms (n=49) (**Fig. S27-30, Supplemental Notes**). Here, the challenge will be not only to finish these regions but to represent changes in meaningful ways such that ectopic exchange events among acrocentric short arms [Garrison et al. companion], interlocus gene conversion among SDs (Vollger, DeWitt, et al. 2022), hypermutability and saltatory amplifications in satellite DNA (Logsdon et al. 2021; Altemose et al. 2022) can be adequately captured. Alternate graph-based approaches, such as PGGB, hold tremendous promise in this regard, but true representation of such diversity requires a fundamental understanding of the mutational processes that have shaped these regions. Complete sequence and characterization of these more complex mutational processes both from a population and familial level (Noyes et al. 2022; Vollger, Guitart, et al. 2022) will facilitate the development of more sophisticated and more representative pangenome graphs in the future.

## METHODS

### Set of evaluated *de novo* assemblies

*De novo* assemblies evaluated in this study have been obtained from two different sources that are part of two international consortia: HGSVC and HPRC. For HGSVC data, we evaluated a panel of 35 samples of diverse ancestry (AFR - 11, AMR - 5, EUR - 7, EAS - 7, SAS - 5). Of those there are 30 and 14 samples with CLR and CSS PacBio data, respectively (9 samples - 3 trios - have available both CLR and CCS data). In the HPRC assembly collection, there are 47 samples of mostly African and American ancestry (AFR - 24, AMR - 16, EUR - 1, EAS - 5, SAS - 1) sequenced using CCS PacBio only.

### Alignment of *de novo* assemblies to the reference genome

#### Alignments used for simple contig end evaluation

All *de novo* assemblies have been aligned to the most complete version of the human reference genome T2T-CHM13 (version 1.1) using minimap2 (version 2.22.0) with the following command:

~~~
minimap2 -K 8G -t {threads} -ax asm20 \
     --secondary=no --eqx -s 25000 \
     {input.ref} {input.query} \
     | samtools view -F 4 -b - > {output.bam}
~~~

Minimap2 had a known issue where some inversions were missed if they were part of another alignment. To alleviate this issue, we realigned the assemblies with the same parameters after hard masking the reference and query to remove regions that were already aligned in the first alignment step. A complete pipeline for this reference alignment is available at: https://github.com/mrvollger/asm-to-reference-alignment

#### Alignments used for contig alignment ends evaluation

All *de novo* assemblies have been aligned to the most complete version of the human reference genome T2T-CHM13 (version 1.1) using a newer minimap2 version (version 2.24.0) with the following command:

~~~
minimap2 -K 8G -t {threads} -x asm20 \
     --secondary=no --eqx -s 25000 \
     {input.ref} {input.query} \
     | samtools view -F 4 -b - > {output.bam}
~~~

A complete pipeline for this reference alignment is available at: https://github.com/mrvollger/asm-to-reference-alignment

### Evaluation of simple contig ends

Contig ends are defined at the first and last aligned base for each contig in the HPRC haplotype-phased assemblies. Alignments were performed as described above, and the terminal position of each contig was determined using rustybam liftover (https://github.com/mrvollger/rustybam). A complete pipeline for identifying contigs ends is included in: https://github.com/mrvollger/asm-to-reference-alignment

### Reading in minimap alignments

Minimap alignments reported in PAF format have been loaded in a set of genomic ranges using custom R function ‘paf2ranges’ with given parameters [min.mapq = 10, min.aln.width = 1000, min.ctg.size = 100000, report.ctg.ends = TRUE, min.ctg.ends = 50000]. At this step we kept alignments with mapping quality equal to or more than 10 and of minimal size 1 kbp. Also contigs with a total size less than 100 kbp have been filtered out.

#### Evaluation of contig alignments ends

After loading all minimap alignments, we extracted terminal contig alignments of size at least 50 kbp. When a total alignment size of a contig to the reference was larger than 5% of an actual contig size, we split such contig into more than one alignment with its own alignment ends. Such splits occur in situations where the end of the contig maps to distal SD pairs or maps across the centromere, thus increasing the mapped contig size in respect to real contig size.

#### Defining genomic region between contig ends and discontinuities within each contig

With minimap alignments loaded in a set of genomic ranges, we set out to determine genomic regions spanning between them. For this we used a custom R function (‘reportGaps’) in order to report genomic ranges between subsequent contig end mappings.

### Strand-seq data generation and data processing

Strand-seq data were generated as follows. EBV-transformed lymphoblastoid cell lines from the 1KG (1000 Genomes Project Consortium et al. 2015) (Coriell Institute) were cultured in BrdU (100 uM final concentration; Sigma, B9285) for 18 or 24 hours, and single isolated nuclei (0.1% NP-40 substitute lysis buffer (Sanders et al. 2017)) were sorted into 96-well plates using the BD FACSMelody cell sorter. In each sorted plate, 94 single cells plus one 100-cell positive control and one 0-cell negative control were deposited. Strand-specific DNA sequencing libraries were generated using the previously described Strand-seq protocol (Falconer et al. 2012; Sanders et al. 2017) and automated on the Beckman Coulter Biomek FX P liquid handling robotic system (Sanders et al. 2019). Following 15 rounds of PCR amplification, 288 individually barcoded libraries (amounting to three 96-well plates) were pooled for sequencing on the Illumina NextSeq500 platform (MID-mode, 75 bp paired-end protocol).

The demultiplexed FASTQ files were aligned to the T2T-CHM13 (v1.1) reference assembly using BWA aligner (version 0.7.17-r1188) and SAMtools (version 1.10). Duplicate reads were marked using sambamba (version 1.0). Low-quality libraries were excluded from future analyses if they showed low read counts, uneven coverage, or an excess of ‘background reads’ yielding noisy single-cell data, as previously described (Porubský et al. 2016; Sanders et al. 2017). Aligned BAM files were used for assembly evaluations as described below.

### Evaluation of assembly quality using Strand-seq

For a set of eight HPRC samples (HG01123, HG01258, HG01358, HG01361, HG01891, HG02257, HG02486, HG02559) for which corresponding Strand-seq data are available, we evaluated the directional and structural contiguity of such assemblies.

#### Evaluation of misorientations or unresolved homozygous inversions

To evaluate any changes in orientation we first processed each selected Strand-seq library using breakpointR with the following parameters: windowsize = 2000000, binMethod = ‘size’, pairedEndReads = TRUE, min.mapq = 10, genoT = ‘binom’, background = 0.1, minReads = 100. Next, we created so-called ‘composite files’ that concatenate directional reads across all libraries using breakpointR function ‘synchronizeReadDir’. We set to detect any changes in directionality by running breakpointR on such composite files with the following parameters: windowsize = 10000, binMethod = “size”, pairedEndReads = FALSE, genoT = ‘binom’, background = 0.1, peakTh = 0.25, minReads = 50. Misorientation and unresolved homozygous inversions are reported as regions that genotypes as having the majority of minus oriented reads (‘ww’, Watson-Watson strand state) while one would expect all Strand-seq reads to map in plus orientation (‘cc’, Crick-Crick strand state) if the assembly is correctly oriented throughout each contig.

#### Evaluation of phasing accuracy for selected PGAS assemblies

We evaluated phasing accuracy for HPRC samples (HG01123, HG01258, HG01358, HG01361, HG01891, HG02257, HG02486, HG02559) for which corresponding Strand-seq data are available and thus both HPRC and PGAS assemblies could be produced. In this analysis we consider trio-based HPRC assemblies as the gold standard for phasing evaluation. We used PAV (v1.1.2) to call SNVs in phased HPRC assemblies as described previously (Ebert et al. 2021). To search for large-scale switch errors, we split phased PGAS assemblies into 1 Mbp long chunks. Subsequently, we used whatshap (version 1.0) to assign each 1 Mbp chunk to either haplotype 1 or 2 based on a trio-based set of phased SNVs. For each sample we evaluated a fraction of wrongly assigned 1 Mbp segments separately for haplotype 1 and 2 across all autosomes. Visually we detected two large-scale switch errors on chromosome 9 in sample HG01891. There was one switch error around position 42 Mbp near centromere and the other near the end of chromosome 9 at position 137.3 Mbp.

#### Evaluation of inversion resolution for selected PGAS assemblies

In order to evaluate performance of trio-based and trio-free assemblies to resolve inversion, we selected a set of large inversions (≥100 kbp) from the previous study (Porubsky et al. 2022). We mapped inversion coordinates from GRCh38 space to T2T-CHM13 (v1.1) coordinates using minimap2 (version 2.20) using following parameters: --secondary=no --eqx -ax asm20 -r 100,1k -z 10000,50. We selected a set of 20 inverted sites (≥100 kbp) with a clear Strand-seq inversion pattern. For dotplot visualization purposes we added extra padding on each side of the inversion equal to the size of the inversion but no less than 2 Mbp. We extracted assembly alignments to the reference T2T-CHM13 (v1.1) corresponding to these regions from each trio-based and trio-free phased assembly using rustybam (version 0.1.27) function ‘liftover’. Next, we exported a FASTA file from each assembly based on subsetted region-specific paf files. We used nucmer (mummer v3.23) with the following parameters: --mum --coords, to align each fasta file to the reference sequence (T2T-CHM13 v1.1). We visualized alignments for each assembly in each inverted region as dotplot. Each dotplot was evaluated manually. Inversion was deemed to be resolved if an inversion can be traced in a single contig in both haplotype and if the inversion status in both haplotypes matches reported inversion genotype presented in (Porubsky et al. 2022).

### Definition of centromeric satellite DNA

In this study centromeric satellite DNA was defined based on T2T-CHM13 annotation obtained from UCSC Table Browser. Annotation was obtained for T2T-CHM13 (v1.1) reference from annotation group ‘Centromeres and Telomeres’ and annotation track ‘CenSat Annotation’. We define centromeric satellite DNA as regions annotated as ‘hsat’ (human-satellites), ‘bsat’ (beta-satellites) and ‘hor’ (alpha-satellites HOR array).

### Protein-coding genes annotation

Gene annotation used in this study is based on T2T-CHM13 annotation obtained from UCSC Table Browser. Annotation was obtained for T2T-CHM13 (v1.1) reference from annotation group ‘Genes’ and annotation track ‘CAT Genes + LiftOff V4’. When reporting gene overlap, we selected only protein-coding genes. Any CHM13-specific genes were not considered. Lastly, subsequent ranges of the same gene were collapsed.

### Evaluation of ONT alignments

Available ONT reads (**Data Availability**) for 33 HPRC samples were aligned to the T2T-CHM13 (version 1.1) reference assembly using minimap2 (version 2.24) and filtered secondary alignments using samtools (version 1.9). We run the alignments with the following parameters:

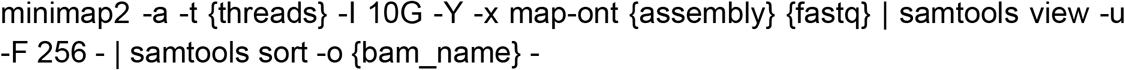

Obtained alignments were exported as read alignment positions in BED format. Only reads with mapping quality 10 and more were retained for further analysis. We tested each reported assembly gap region per sample and per haplotype if such region is spanned by 10 and more ONT reads to assume that such assembly gap could eventually be closed by underlying ONT reads.

### Dinucleotide frequency in frequent assembly breaks

Out of the total of 592 defined frequent assembly breaks, we extracted 44 regions where there is an assembly break in half and more HPRC assemblies. Next, we extracted the T2T-CHM13 FASTA sequence corresponding to these regions (n=44). We calculated the total number of three dinucleotides (TA, TC and GA) in nonoverlapping 100 bp long sequence chunks (bins). To define dinucleotide enriched bins, we transformed binned dinucleotide counts into the z-scores and marked bins with z-score ≥1.96 (95% confidence interval) as dinucleotide enriched. The size of dinucleotide tracts have been estimated as the number of enriched bins * 100 (bin size).

We also investigated FASTA sequence from the previously defined regions (n=44) in all HPRC assemblies along with a number (n=18) of nonhuman primates assemblies. We processed only those assemblies that span defined regions in a single contig and map to defined breakpoints in T2T-CHM13 coordinates (+/-100 bp). Next we transformed observed dinucleotide counts into z-scores as outlined above. Based on visual inspection, we selected 27/44 regions with observable differences in size of dinucleotide tracts between human and nonhuman primate assemblies (**Table S5**).

### Defining regions of putative structural variation

We examined large contig alignment discontinuities as gaps within a single contig alignment that are smaller than 1 Mbp. We classified a contig alignment discontinuity as a ‘contraction’ if the alignment gap (in target sequence coordinates) is larger than the corresponding gap within a contig (in query sequence coordinates) (**Fig. 1A, iii**). In contrast, we classified a contig alignment discontinuity as an ‘expansion’ if the alignment gap (in target sequence coordinates) is smaller than the corresponding gap within a contig (in query sequence coordinates). The number of unaligned bases is defined as the size of the gap in query sequence coordinates. Predicted size of the contraction and expansions was defined as a difference in size between gap in target and query coordinates. We marked contig alignment discontinuities that are within or close (+/-1 Mbp) to centromeric satellite DNA (marked as ‘CENSAT’). This is because contig assemblies and alignments within and nearby centromeres are complicated by the repetitive nature of centromeric satellites and high degree of SDs in these regions. We summarized predicted sites of contraction and expansion into a set of nonredundant regions constructed from sites where contraction and expansion is observed in at least five assemblies and predicted event size is 100 bp and longer (**Table S6**).

### Detection of CNV regions

In order to define regions that are likely copy number variable in any given sample, we searched for regions where there are overlapping contig alignments in respect to the T2T-CHM13 (v1.1) reference. In this analysis we considered only autosomes and we filtered out regions that overlap centromeric satellites. We opt to validate putative CNV regions using short-read-based copy number profiles obtained for 44/47 HPRC samples. Short Illumina reads were computationally parsed into 36 bp segments and aligned to a hardmasked T2T-CHM13 (v1.1) reference using mrsFAST (Hach et al., 2010) allowing an edit distance of 2. Read-depth-based copy number estimates were generated using the FastCN (Pendleton et al., 2018) software package, which uses known copy number stable regions to correct for Illumina sequencing GC bias and convert read depth to diploid copy number over windows containing 500 unmasked base–pairs.

Due to the mapping of short-reads to a single paralogous copy in the genome we set to determine sample-specific copy number by establishing reference copy number of paralogous regions in T2T-CHM13 (v1.1). We did this by splitting T2T-CHM13 (v1.1) sequence into the same 36 bp subsequences with a slide of 1 to cover all kmers in the reference. These kmers were mapped back to the reference using mrsFAST and copy number determined via FastCN. This will be referred to as the kmer-ized CHM13 reference copy number.

We defined sample-specific CNV regions as those with diploid copy number less than 10 and at least one diploid copy number increase compared to the kmer-ized CHM13 reference copy number. Sample-specific regions with diploid copy number of 2 and/or no delta from the kmer-ized CHM13 reference copy number were defined as not copy number variable and marked as ‘noCN’. Regions where there is an observable diploid copy number increase but the overall, sample-specific copy number is greater than 10 we marked as ‘more10CN’. Regions that do not fall into any of the above categories were marked as ‘none’.

### Analysis of pangenome brnn regions

Genomic regions that have been excluded from the CHM13-based pangenome graph construction were obtained from https://github.com/human-pangenomics/hpp_pangenome_resources#masked-sequenc. Detailed description of how these regions were defined is reported in the link above. We next took the file ‘hprc-v1.0-mc-chm13.clipped-intervals.bed.gz’ and for each genomic region we extracted the FASTA sequence from a corresponding phased assembly. We next aligned these to T2T-CHM13 (v1.1) reference using minimap2 (version 2.24) with the following parameters: --secondary=no --eqx -ax asm20 -r 100,1k -z 10000,50. Next we kept only alignments of minimum mapping quality of ≥10 and also we excluded any alignments from mitochondrial DNA.

### Generation of DeepVariant SNP calls for false LOH detection

Alignments of raw PacBio HiFi reads (from eight samples: HG02486, HG02572, HG02622, HG02886, HG03516, HG03540, HG03579) to T2T-CHM13 (v1.1) were made with pbmm2 (https://github.com/PacificBiosciences/pbmm2) using the ‘CCS’ preset. DeepVariant calls were generated using DeepVariant (Poplin et al. 2018) version 1.4.0 and the ‘PACBIO’ pretrained model.

## DATA AND CODE AVAILABILITY

PacBio HiFi, ONT and Strand-seq data have been deposited into NCBI Sequence

Read Archive (SRA) under the following bioproject IDs: PRJNA850430, PRJNA731524 and PRJEB54100.

The T2T-CHM13 (v1.1) assembly can be found on NCBI (GCA_009914755.3).

DeepVariant callset for selected samples (HG02486, HG02572, HG02622, HG02886, HG03516, HG03540, HG03579) is available at Zenodo DOI: 10.5281/zenodo.6762544

Dipcall callset for selected samples (HG02486, HG02572, HG02622, HG02886, HG03516, HG03540, HG03579) was obtained from: https://s3-us-west-2.amazonaws.com/human-pangenomics/index.html?prefix=working/HPRC/{sample.id}/assemblies/year1_f1_assembly_v2_genbank/assembly_qc/dipcall/{sample.id}.f1_assembly_v2_genbank.dip.vcf.gz

Download location of ONT data used in this study are reported in **Table S9**.

Download location of Strand-seq data for eight samples (HG01123, HG01258, HG01358, HG01361, HG01891, HG02257, HG02486, HG02559) are reported in **Table S9**.

Custom R scripts used in this study to process assembly alignments and report assembly gaps can be obtained from Zenodo data repository, DOI: 10.5281/zenodo.6762544

FASTA sequences from selected low complexity regions (n=27) can be obtained from Zenodo data repository, DOI: 10.5281/zenodo.6762544

## AUTHOR CONTRIBUTIONS

Conceptualization and design: D.P., M.R.V and E.E.E.; Assembly gap, inversion resolution and structurally complex region analysis D.P.; Contig end enrichment analysis: M.R.V.; Production of HGSVC assemblies: P.E. and T.M.; Production of HPRC assemblies: HPRC group; Strand-seq data generation: P.H., A.S.D., C.S. and J.O.K.; Bioinformatics support: W.T.H and A.N.R.; Organization of tables and supplementary material: D.P.; Display items: D.P. and M.R.V.; Resources: HPRC, G.H., B.P. and E.E.E.; Manuscript writing: D.P., M.R.V., and E.E.E., with input from all authors.

## ACKNOWLEDGEMENTS

We thank Tonia Brown for assistance in editing this manuscript. T.M. and P.E. acknowledge the support and computational infrastructure provided by the Centre for Information and Media Technology at Heinrich Heine University Düsseldorf.

This article is subject to HHMI’s Open Access to Publications policy. HHMI lab heads have previously granted a nonexclusive CC BY 4.0 license to the public and a sublicensable license to HHMI in their research articles. Pursuant to those licenses, the author-accepted manuscript of this article can be made freely available under a CC BY 4.0 license immediately upon publication.

## Human Pangenome Reference Consortium (HPRC)

Haley J. Abel^1^, Lucinda L Antonacci-Fulton^2^, Mobin Asri^3^, Gunjan Baid^4^, Anastasiya Belyaeva^4^, Konstantinos Billis^5^, Guillaume Bourque^6,7,8^, Silvia Buonaiuto^9^, Andrew Carroll^4^, Mark JP Chaisson^10^, Pi-Chuan Chang^4^, Xian H. Chang^3^, Haoyu Cheng^11,12^, Justin Chu^11^, Sarah Cody^2^, Vincenza Colonna^9,13^, Daniel E. Cook^4^, Omar E. Cornejo^14^, Mark Diekhans^3^, Daniel Doerr^15^, Peter Ebert^15^, Jana Ebler^15^, Evan E. Eichler^16,17^, Jordan M. Eizenga^3^, Susan Fairley^5^, Olivier Fedrigo^18^, Adam L. Felsenfeld^19^, Xiaowen Feng^11,12^, Christian Fischer^20^, Paul Flicek^5^, Giulio Formenti^18^, Adam Frankish^5^, Robert S. Fulton^2^, Yan Gao^21^, Shilpa Garg^22^, Erik Garrison^13^, Carlos Garcia Giron^5^, Richard E. Green^23,24^, Cristian Groza^25^, Andrea Guarracino^26^, Leanne Haggerty^5^, Ira Hall^27,28^, William T Harvey^16^, Marina Haukness^3^, David Haussler^3,17^, Simon Heumos^29,30^, Glenn Hickey^3^, Thibaut Hourlier^5^, Kerstin Howe^31^, Miten Jain^32^, Erich D. Jarvis^33,17^, Hanlee P. Ji^34^, Alexey Kolesnikov^4^, Jan O. Korbel^35^, HoJoon Lee^34^, Alexandra P. Lewis^16^, Heng Li^11,12^, Wen-Wei Liao^2,36,27^, Shuangjia Lu^27^, Tsung-Yu Lu^10^, Julian K. Lucas^3^, Hugo Magalhães^15^, Santiago Marco-Sola^37,38^, Pierre Marijon^15^, Tobias Marschall^15^, Fergal J. Martin^5^, Jennifer McDaniel^39^, Karen H. Miga^3^, Matthew W. Mitchell^40^, Jean Monlong^3^, Jacquelyn Mountcastle^18^, Katherine M. Munson^16^, Moses Njagi Mwaniki^41^, Maria Nattestad^4^, Adam M. Novak^3^, Hugh E. Olsen^3^, Nathan D. Olson^39^, Benedict Paten^3^, Trevor Pesout^3^, Adam M. Phillippy^42^, David Porubsky^16^, Pjotr Prins^13^, Daniela Puiu^43^, Allison A Regier^2^, Arang Rhie^42^, Samuel Sacco^44^, Ashley D. Sanders^45^, Valerie A. Schneider^46^, Baergen I. Schultz^19^, Kishwar Shafin^4^, Jonas A. Sibbesen^47^, Jouni Sirén^3^, Michael W. Smith^19^, Heidi J. Sofia^19^, Ahmad N. Abou Tayoun^48,49^, Françoise Thibaud-Nissen^46^, Chad Tomlinson^2^, Francesca Floriana Tricomi^5^, Flavia Villani^13^, Mitchell R. Vollger^16,50^, Justin Wagner^39^, Ting Wang^51^, Jonathan M. D. Wood^31^, Aleksey V. Zimin^43,52^, Justin M. Zook^39^

1. Division of Oncology, Department of Internal Medicine, Washington University School of Medicine, St. Louis, MO 63110, USA
2. McDonnell Genome Institute, Washington University School of Medicine, St. Louis, MO 63108, USA
3. UC Santa Cruz Genomics Institute, University of California, Santa Cruz, 1156 High St, Santa Cruz, CA, USA
4. Google LLC, 1600 Amphitheater Pkwy, Mountain View, CA 94043, USA
5. European Molecular Biology Laboratory, European Bioinformatics Institute, Wellcome Genome Campus, Cambridge, CB10 1SD, UK
6. Department of Human Genetics, McGill University, Montreal, Québec H3A 0C7, Canada
7. Canadian Center for Computational Genomics, McGill University, Montreal, Québec H3A 0G1, Canada
8. Institute for the Advanced Study of Human Biology (WPI-ASHBi), Kyoto University, Kyoto 606-8501, Japan
9. Institute of Genetics and Biophysics, National Research Council, Naples 80111, Italy
10. University of Southern California, Quantitative and Computational Biology, 3551 Trousdale, Pkwy, Los Angeles, CA, USA
11. Department of Data Sciences, Dana-Farber Cancer Institute, Boston, MA 02215, USA
12. Department of Biomedical Informatics, Harvard Medical School, Boston, MA 02215, USA
13. Department of Genetics, Genomics and Informatics, University of Tennessee Health Science Center, Memphis, TN 38163, USA
14. School of Biological Sciences, Washington State University, Pullman WA 99163, USA
15. Institute for Medical Biometry and Bioinformatics, Medical Faculty, Heinrich Heine University Düsseldorf, Düsseldorf, Germany
16. Department of Genome Sciences, University of Washington School of Medicine, Seattle, WA 98195, USA
17. Howard Hughes Medical Institute, Chevy Chase, MD 20815, USA
18. The Vertebrate Genome Laboratory, The Rockefeller University, New York, NY 10065, USA
19. National Institutes of Health (NIH)–National Human Genome Research Institute, Bethesda, MD, USA
20. USA University of Tennessee Health Science Center, Memphis, TN 38163, USA
21. Center for Computational and Genomic Medicine, The Children’s Hospital of Philadelphia, Philadelphia, PA 19104, USA.
22. Department of Biology, University of Copenhagen, Denmark
23. Department of Biomolecular Engineering, University of California, Santa Cruz, 1156 High St., Santa Cruz, CA 95064, USA
24. Dovetail Genomics, Scotts Valley, CA 95066, USA
25. Quantitative Life Sciences, McGill University, Montreal, Québec H3A 0C7, Canada
26. Genomics Research Centre, Human Technopole, Milan 20157, Italy
27. Department of Genetics, Yale University School of Medicine, New Haven, CT 06510, USA
28. Center for Genomic Health, Yale University School of Medicine, New Haven, CT 06510, USA
29. Quantitative Biology Center (QBiC), University of Tübingen, Tübingen 72076, Germany
30. Biomedical Data Science, Department of Computer Science, University of Tübingen, Tübingen 72076, Germany
31. Tree of Life, Wellcome Sanger Institute, Hinxton, Cambridge, CB10 1SA, UK
32. Northeastern University, Boston, MA 02115, USA
33. The Rockefeller University, New York, NY 10065, USA
34. Division of Oncology, Department of Medicine, Stanford University School of Medicine, Stanford, CA, 94305, USA
35. European Molecular Biology Laboratory, Genome Biology Unit, Meyerhofstr. 1, 69117 Heidelberg, Germany
36. Department of Medicine, Washington University School of Medicine, St. Louis, MO 63110, USA
37. Computer Sciences Department, Barcelona Supercomputing Center, Barcelona, Spain
38. Departament d’Arquitectura de Computadors i Sistemes Operatius, Universitat Autònoma de Barcelona, Barcelona, Spain
39. Material Measurement Laboratory, National Institute of Standards and Technology, Gaithersburg, MD 20877, USA
40. Coriell Institute for Medical Research, Camden, NJ 08103, USA
41. Department of Computer Science, University of Pisa, Pisa 56127, Italy
42. Genome Informatics Section, Computational and Statistical Genomics Branch, National Human Genome Research Institute, National Institutes of Health, Bethesda, MD 20892, USA
43. Department of Biomedical Engineering, Johns Hopkins University, Baltimore 21218, MD, USA
44. Department of Ecology & Evolutionary Biology, University of California, Santa Cruz, 1156 High St, Santa Cruz, CA, USA
45. Berlin Institute for Medical Systems Biology, Max Delbrück Center for Molecular Medicine in the Helmholtz Association, Berlin, Germany
46. National Center for Biotechnology Information, National Library of Medicine, National Institutes of Health, Bethesda, MD 20894, USA
47. Center for Health Data Science, University of Copenhagen, Denmark
48. Al Jalila Genomics Center of Excellence, Al Jalila Children’s Specialty Hospital, Dubai, UAE
49. Center for Genomic Discovery, Mohammed Bin Rashid University of Medicine and Health Sciences, Dubai, UAE
50. Division of Medical Genetics, University of Washington School of Medicine, Seattle, WA 98195, USA
51. Department of Genetics, Washington University School of Medicine, St. Louis, MO 63110, USA
52. Center for Computational Biology, Johns Hopkins University, Baltimore, MD 21218, USA

