## Supplemental Figures and Notes for "Gaps and complex structurally variant loci in phased genome assemblies"

### SUPPLEMENTAL FIGURES (1-30):

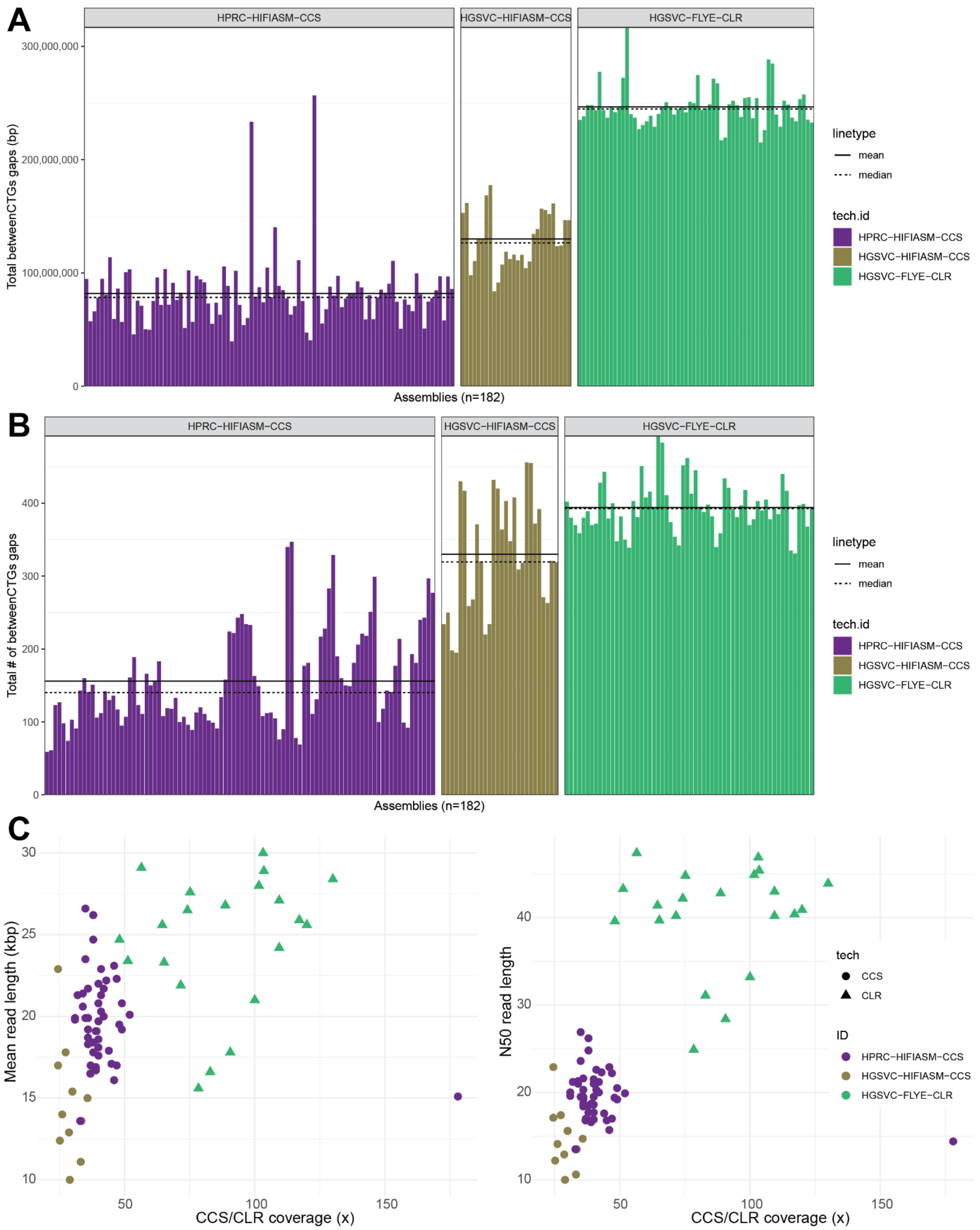


**Figure S1: Number of assembly breaks per haploid genome assembly.**

**A**) A total number of base pairs residing in between any two different contig alignment ends stratified by assembly technology (HGSVC-FLYE-CLR (n=60), HGSVC-PEREG-CCS (n=28), HGSVC-HIFIASM-CCS (n=28), HPRC-HIFIASM-CCS (n=94)). Mean and median values are highlighted as solid and dashed horizontal lines, respectively. **B**) The total number of assembly breaks defined by gap in between two different contig alignment ends stratified by assembly technology. Mean and median values are highlighted as solid and dashed horizontal lines, respectively. **C**) Scatterplot showing circular consensus sequencing (CCS) and continuous long-read (CLR) data QC stratified by assembly technology (point shape) as well as study (Ebert et al. 2021; Liao et al. 2022) and used assembler (see legend ID).

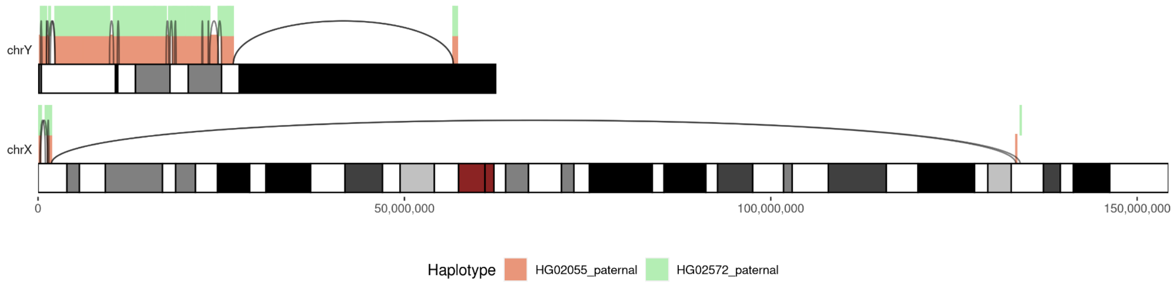


### Figure S2: Unexpected contig mapping to chromosome X in paternal haplotypes.

Chromosome X and Y ideograms showing contig mappings (colored rectangles on top of each ideogram) for two males samples (HG02055, HG02572). Gaps between subsequent contigs are denoted by black lines.

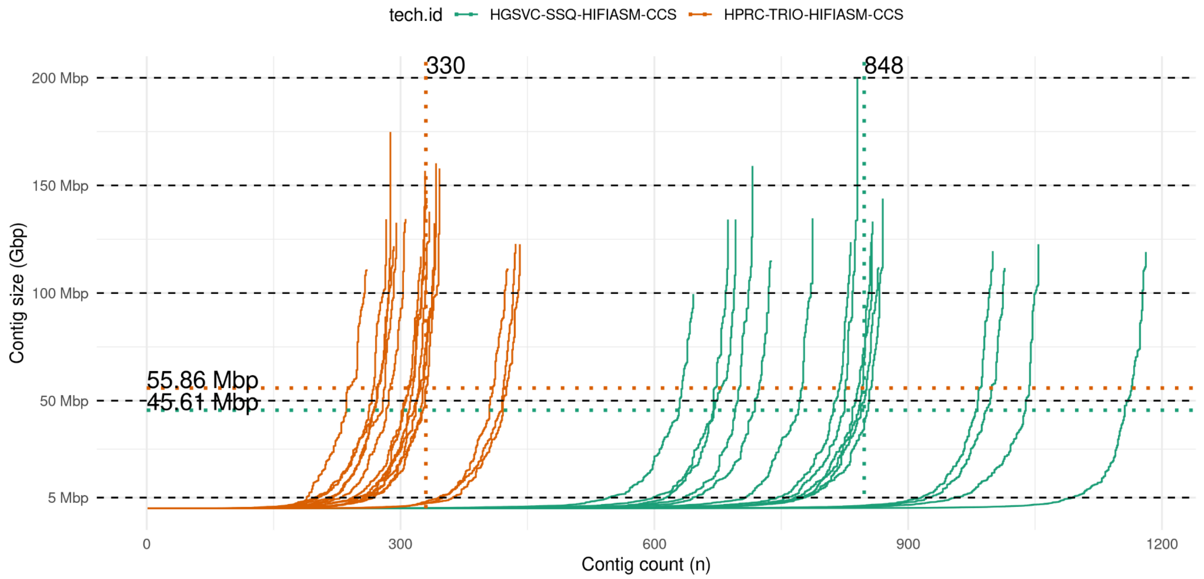


### Figure S3: Evaluation of assembly contiguity between trio-based and trio-free assemblies.

Each line represents a size-ordered set of contigs for 16 phased assemblies (8 human samples) separately for trio-based (orange) and trio-free (green) assembly pipelines. Vertical dotted lines show median number of contigs per assembly pipeline, while horizontal dotted lines shows N50 values per assembly pipeline.

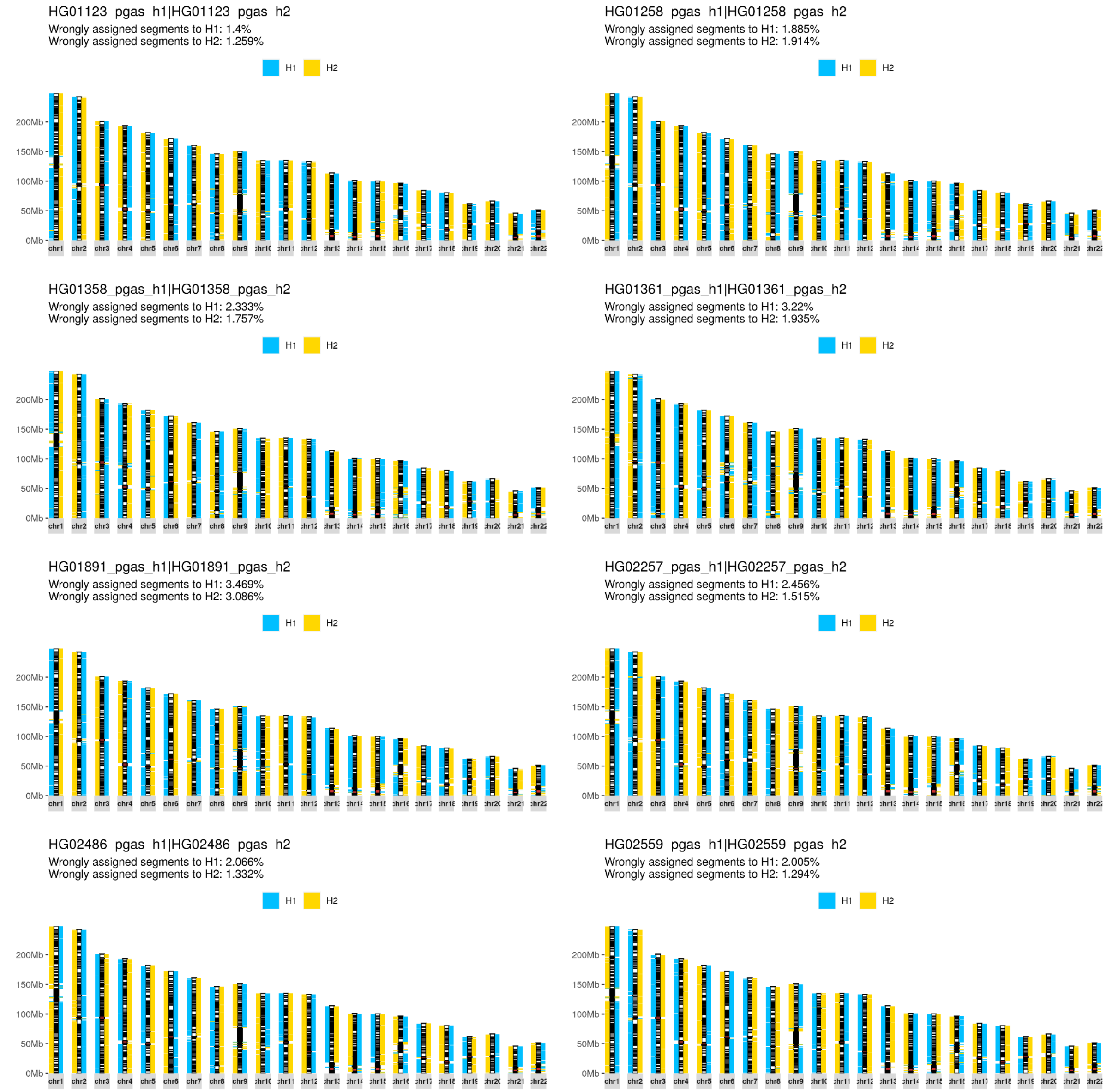


**Figure S4: Evaluation of phasing accuracy of trio-free assemblies.**
Phased trio-free assemblies (n=8) fragmented into 1 Mbp sized blocks (left from ideogram - H1, right from ideogram - H2) are assigned to either haplotype 1 or 2 (H1 - blue, H2 - yellow) using single-nucleotide polymorphims determined in trio-based assemblies with respect to the reference T2T-CHM13 (v1.1) (**Methods**).

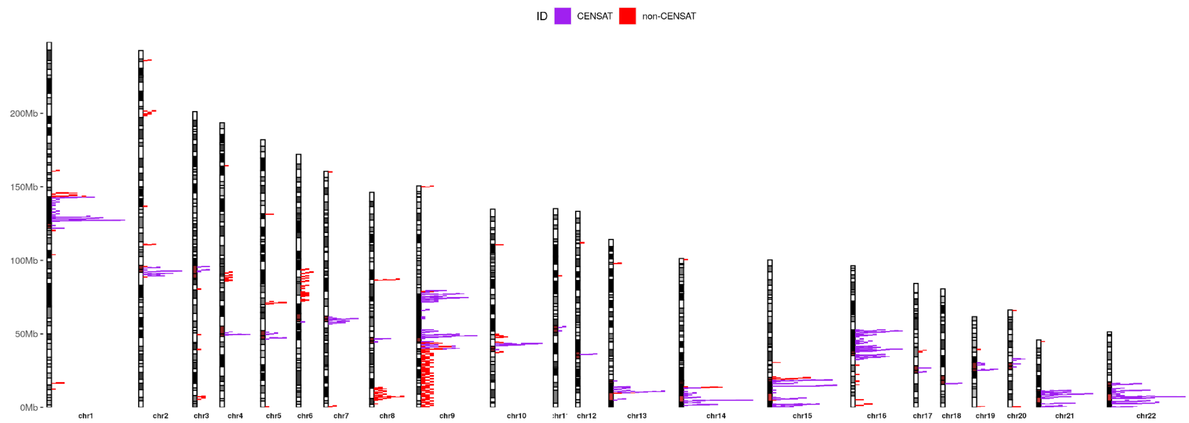


### Figure S5: Genome-wide distribution of wrongly phased segments.

An ideogram showing the position of all 1 Mbp phased assembly segments assigned to a wrong parental haplotype based on trio-based phasing (**Methods**). All wrongly assigned assembly segments are piled up across all evaluated assemblies (n=8, 16 phased assemblies). Each segment is color based on the overlap with centromeric satellites (CENSAT). Note: There is a pileup of wrongly assigned segments on a p-arm of chromosome 9 due to a large-scale phasing error in a single sample (HG01891) (**Fig. 2B**).

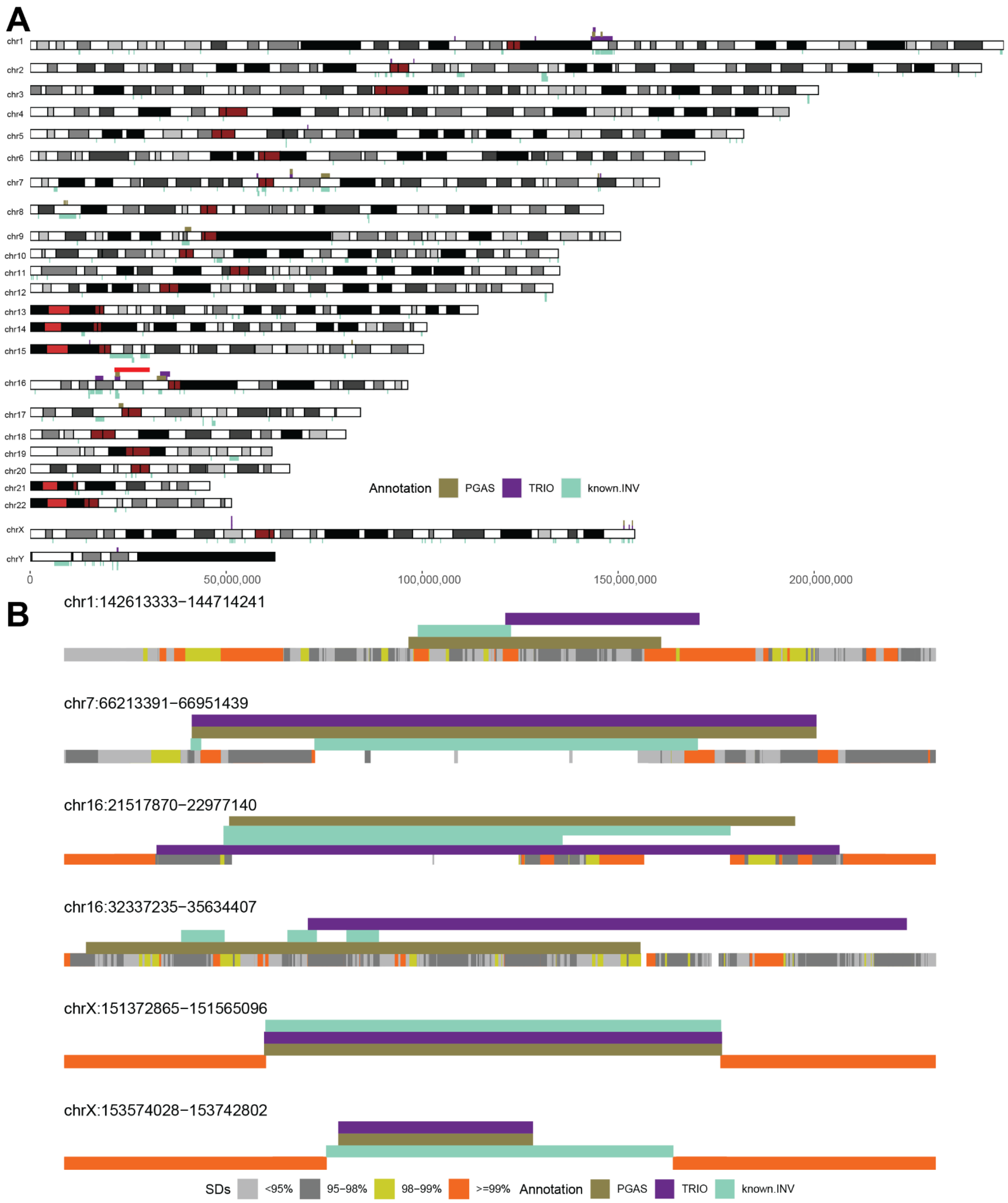


### Figure S6: Wrongly resolved homozygous inversions.

**A**) Genomic positions of wrongly resolved homozygous inversions in trio-based (TRIO, purple, n=22) and trio-free (PGAS, brown, n=15) assemblies above the ideogram. Below the ideogram are the positions of known inverted regions from a recent study (Porubsky et al. 2022). The red rectangle highlights 16p11.2-p12.2 microdeletion/microduplication syndrome with an unresolved homozygous inversion at its proximal breakpoint. **B**) Known inversion sites (known.INV, turquoise) from A that were unresolved by both trio-based (TRIO, purple) and trio-free (PGAS, brown) assembly pipelines. At the bottom of each region there is an SD annotation colored by sequence identity.

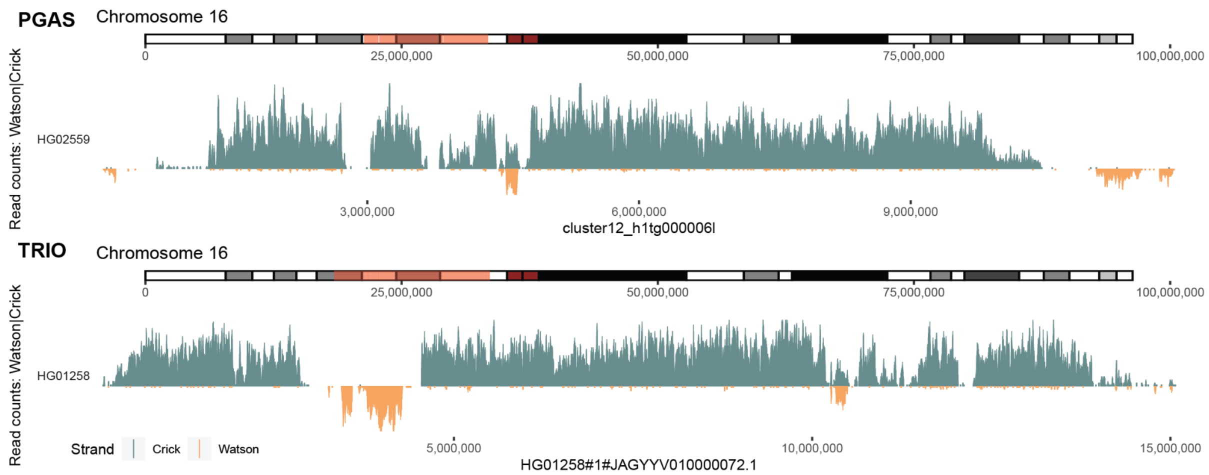


### Figure S7: Examples of unresolved homozygous inversions.

Strand-seq directional read coverage for four selected contigs reported as binned (bin size: 10,000, step size: 1,000) read counts represented as vertical bars above (teal; Crick read counts) and below (orange; Watson read counts) the midline. A region with roughly equal coverage of Watson and Crick reads represents a heterozygous inversion (purple arrowheads) as only one homologue is inverted with respect to the contig assembly. A region with reads aligned only in the Watson orientation points to an unresolved inversion in the contig assembly as both homologs are expected to be inverted. Lastly, regions with only Crick reads reported are those where contig assembly directionality agrees with the Strand-seq data.

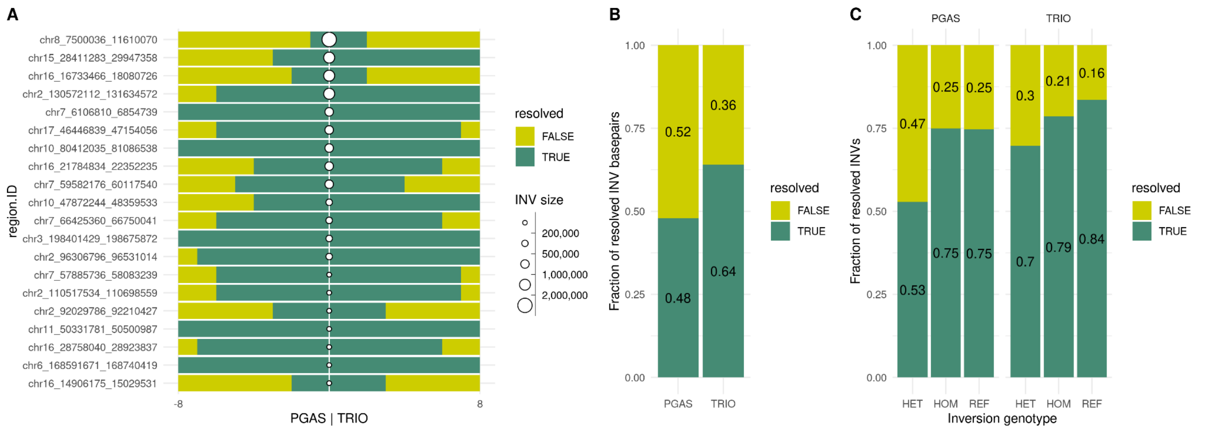


### Figure S8: Evaluation of large inversion resolution between trio-based and trio-free assemblies.

**A**) A bargraph reporting the number of human samples (n=8) that are fully informative on inversion status on both haplotypes (TRUE, dark green). Shown separately for trio-free (PGAS, left) and trio-based (TRIO, right) assemblies and for each evaluated inversion site (n=20) larger than 100 kbp (rows). Inversion sites are sorted by size (from top to bottom). Inversion size is represented by the point size at the vertical midline. **B**) Fraction of tested inverted base pairs that have been properly resolved and thus informative of inversion status (TRUE, dark green). **C**) Fraction of tested inversion sites that are fully informative (TRUE, dark green) as a function of inversion genotype (HET - heterozygous, HOM - homozygous inverted, REF - homozygous reference), reported separately for trio-free (PGAS, left) and trio-based (TRIO, right) assemblies

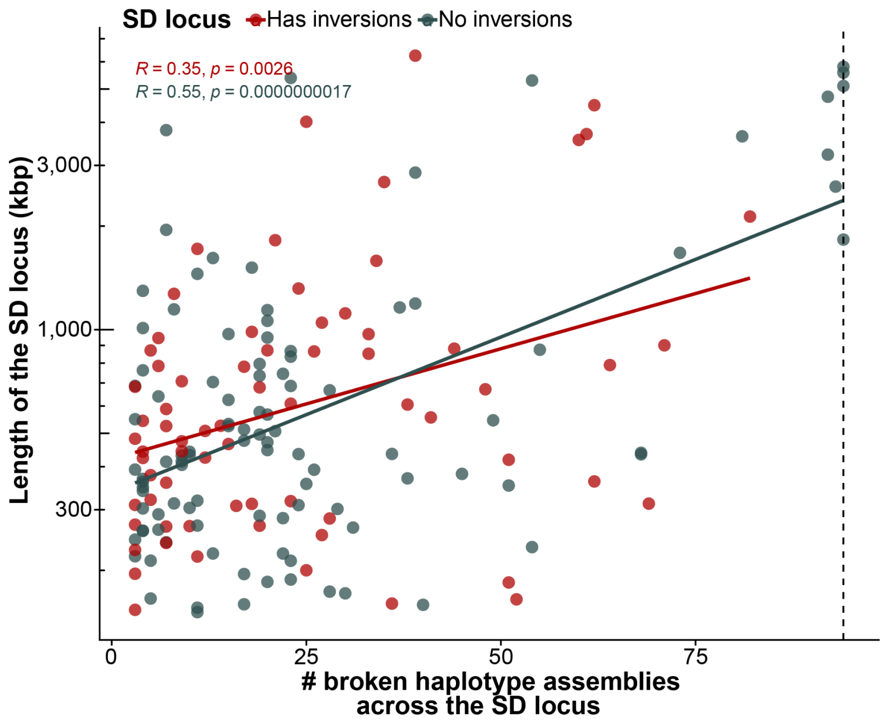


### Figure S9: SD length versus assembly breaks.

Correlation between the length of an SD locus and the number of haplotypes that are contiguously assembled across it. Points are colored by whether the SD locus is polymorphic for inversions (blue) or not (red).

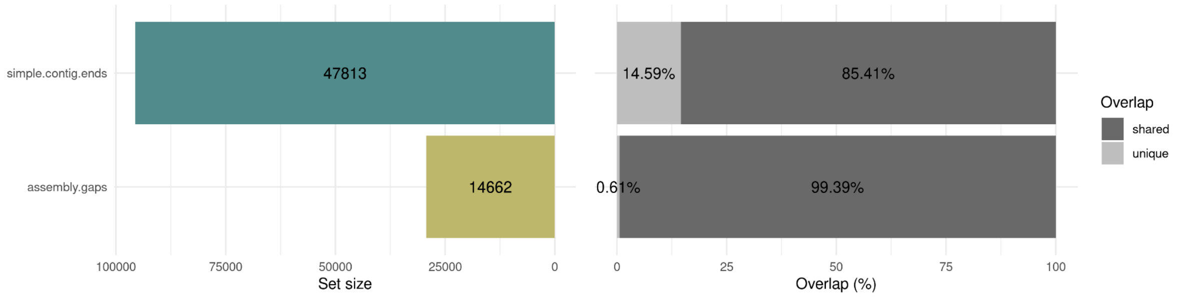


### Figure S10: Comparison between simple contig ends and contig alignment ends.

Overall, we found 85.41% of all simple contig ends fall within +/-10 kbp from gaps defined in between contig alignment ends. Both definitions of assembly breaks agree very well and the small discrepancy is likely caused by small differences in contig end definition and applied filters (**Methods**). For instance, contig alignment ends are not reported for contigs smaller than 100 kbp while simple contig ends are.

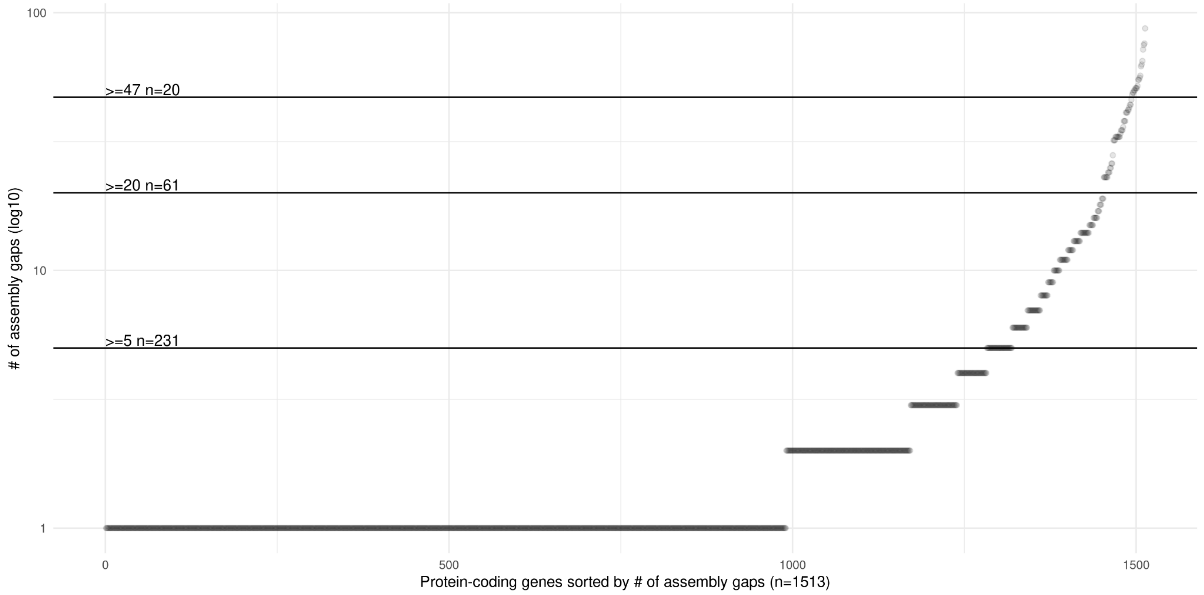


### Figure S11: Protein-coding genes broken at defined assembly gaps.

A scatterplot of all protein-coding genes as they overlap the full set of assembly gaps (n=14,662) across all HPRC assemblies. Horizontal lines highlight subsets of protein-coding genes that overlap with a defined number (y-axis) of assembly gaps and more.

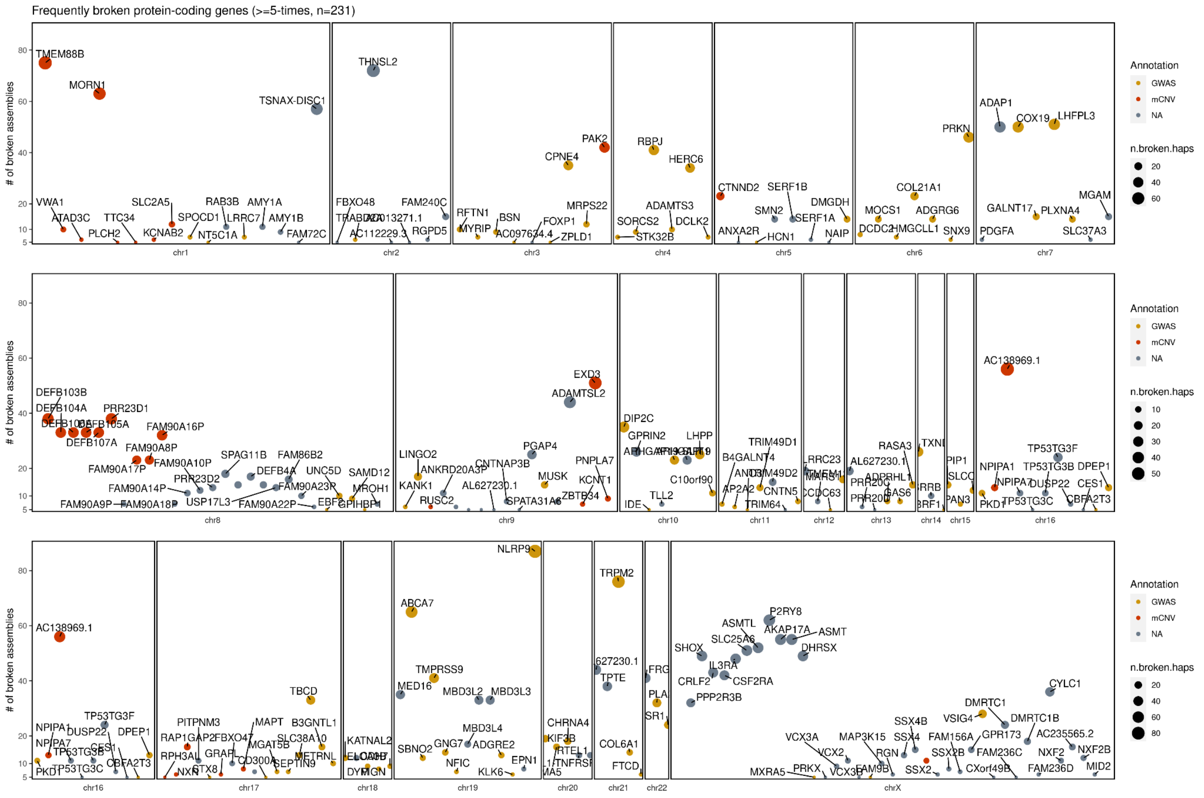


### Figure S12: Protein-coding genes within frequent assembly gaps.

An overview of protein-coding genes (n=231) overlapping with assembly breaks that occurred in five and more assemblies. Each gene is marked by a dot whose size reflects the number of assembly gaps it overlaps. The color of each dot marks if a given gene overlaps with known morbid CNVs or if it was previously reported in GWAS studies (**Table S3**).

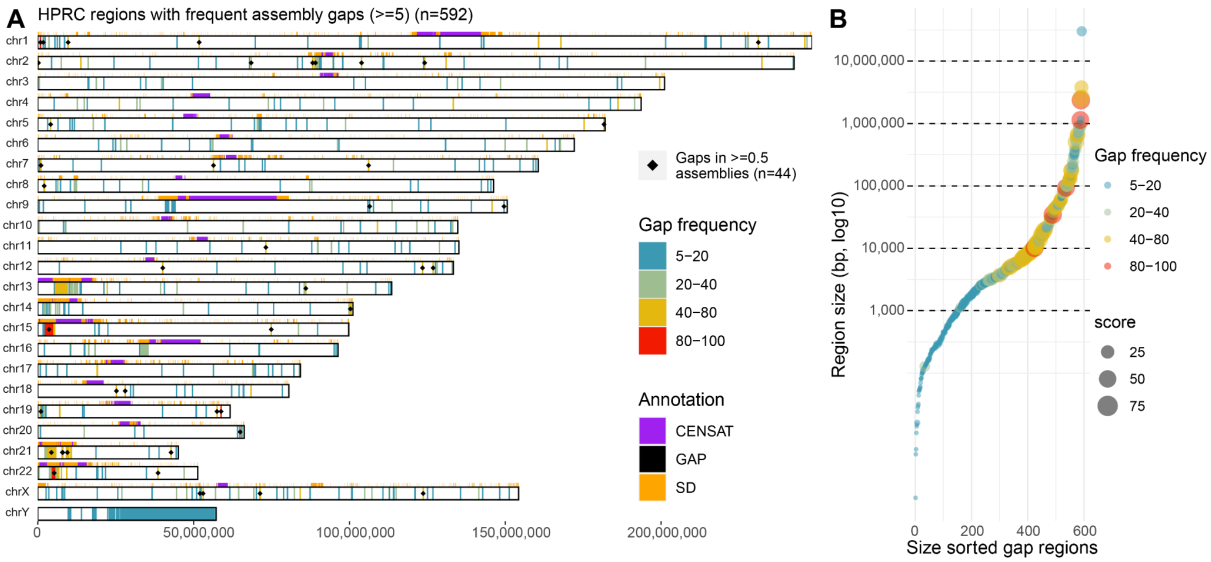


### Figure S13: Frequent assembly gaps.

**A**) Genome-wide distribution or 592 regions where there is an assembly break in at least five different HPRC assemblies (n=94) (**Table S4**). Each genomic region is colored by the number of assembly breaks contained in each region. At the top of each chromosomal bar there is annotation of SDs (orange) and higher order centromeric regions (CENSAT, purple). Regions with assembly break in half and more HPRC assemblies are highlighted by black diamond. **Note:** Some larger regions might contain multiple breaks from a single assembly and thus might exceed the total number of assessed assemblies (n=94). **B**) A scatterplot showing the size distribution of defined frequent assembly breaks ordered on x-axis that are colored (also size of each dot) by the number of assembly breaks defined in each of these regions.

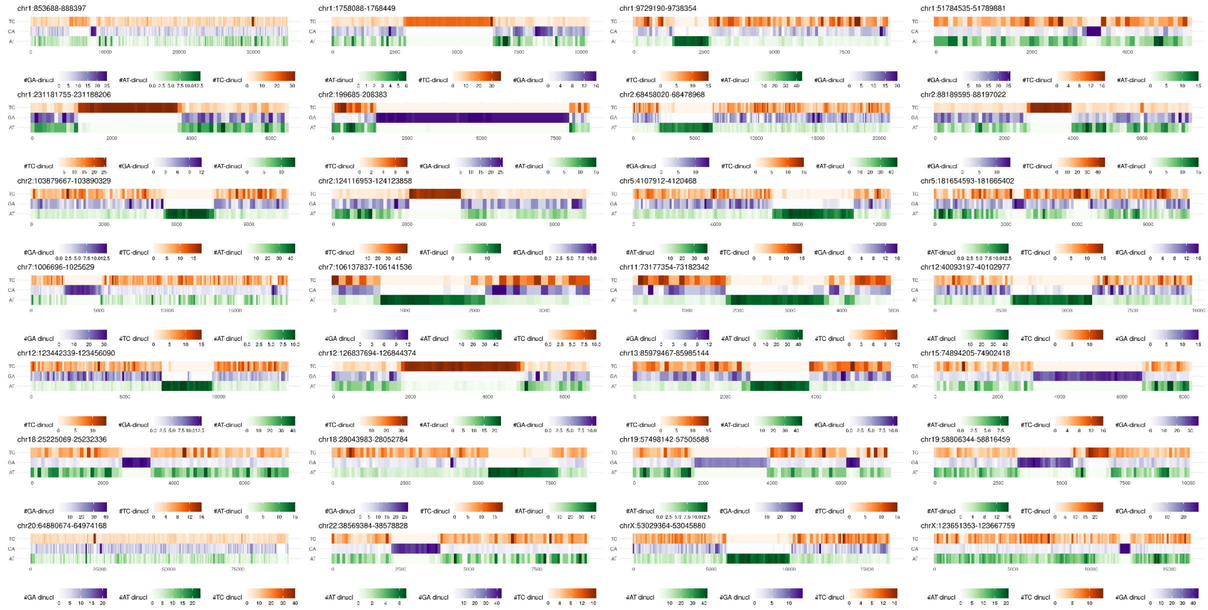

### Figure S14: Frequent assembly gaps in low-complexity regions not associated with SDs.

Visualization of dinucleotide composition of 28 regions where assemblies break frequently (≥0.5 of all HPRC assemblies). Also, none of these regions are associated with SDs. Each region is decomposed into counts of TC, GA, and AT dinucleotides along DNA sequence extracted from T2T-CHM13 (v1.1). Frequency of each dinucleotide is visualized as a heatmap along each region.

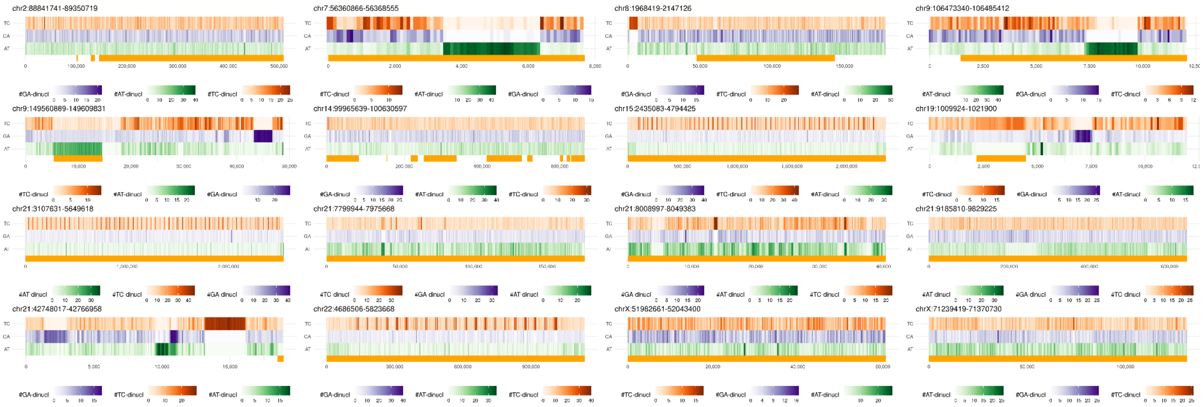

**Figure S15: Frequent assembly gaps associated with SDs.**
Visualization of dinucleotide composition of 16 regions where assemblies break frequently (≥0.5 of all HPRC assemblies). All these regions have positive overlap with known SDs. Each region is decomposed into counts of TC, GA, and AT dinucleotides along DNA sequence extracted from T2T-CHM13 (v1.1). The frequency of each dinucleotide is visualized as a heatmap along each region. Regions that overlap with SDs are highlighted by orange rectangles at the bottom.

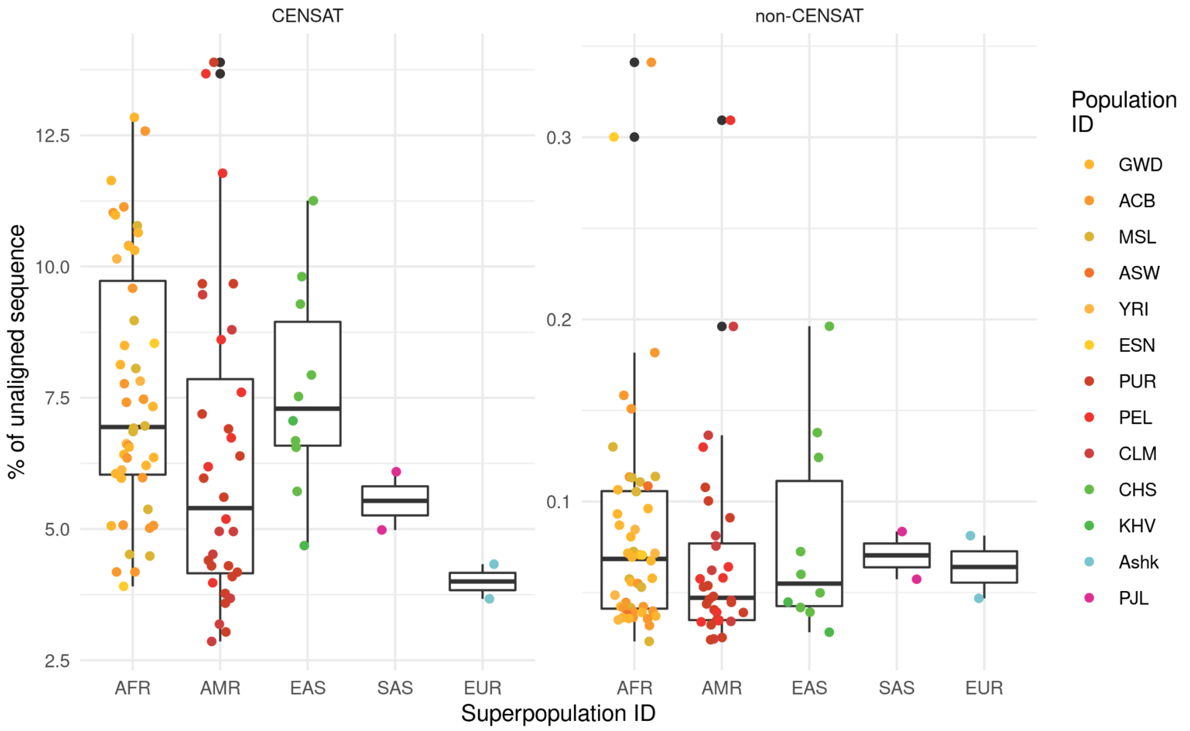


**Figure S16: Extent of assembly bases in contig alignment discontinuities not represented with respect to reference (T2T-CHM13 v1.1).**
Boxplots showing the percentage of contig bases that are either not aligned or are missing with respect to reference genome (T2T-CHM13 v1.1). Each dot represents haploid assembly (n=94) colored by population identifier (see legend) stratified per superpopulation identifier (AFR - African, AMR - American, EAS - East Asian, SAS - South-East Asian and EUR - European) and classification if unassigned contig bases belong to centromeric satellite DNA ('CENSAT') or not ('non-CENSAT').

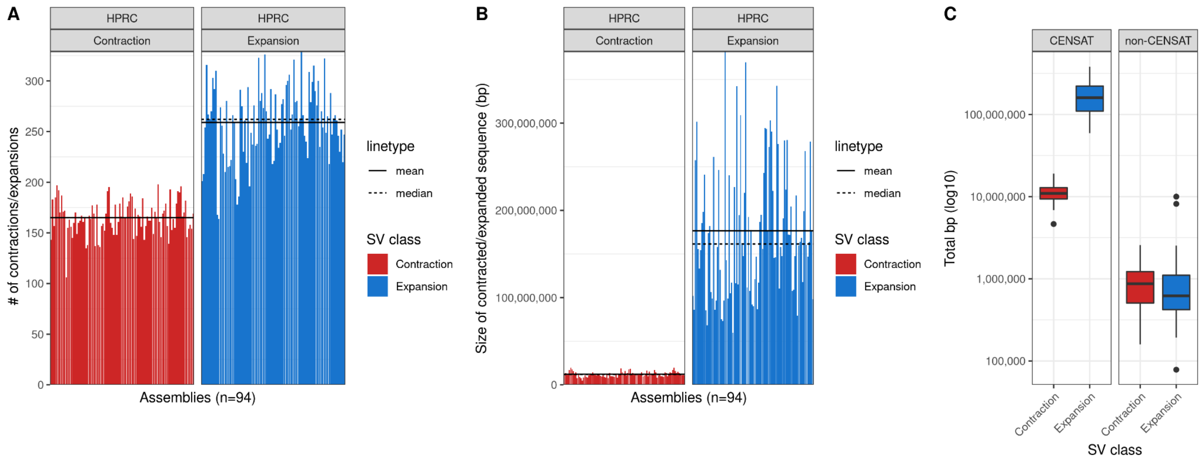


### Figure S17: Extent of contracted and expanded base pairs in discontinuous contig alignments.

**A**) The total number of contig alignment discontinuities stratified by class (red - Contraction, blue - Expansion) across all 94 phased HPRC assemblies. **B**) The total number of base pairs contributed by each class of contig alignment discontinuities (red - Contraction, blue - Expansion) across all 94 phased HPRC assemblies. In both A and B, mean and median values are highlighted as solid and dashed horizontal lines, respectively. **C**) A boxplot showing a proportion of contraction and expansion bases within (CENSAT) and outside (non-CENSAT) of centromeric satellite DNA.

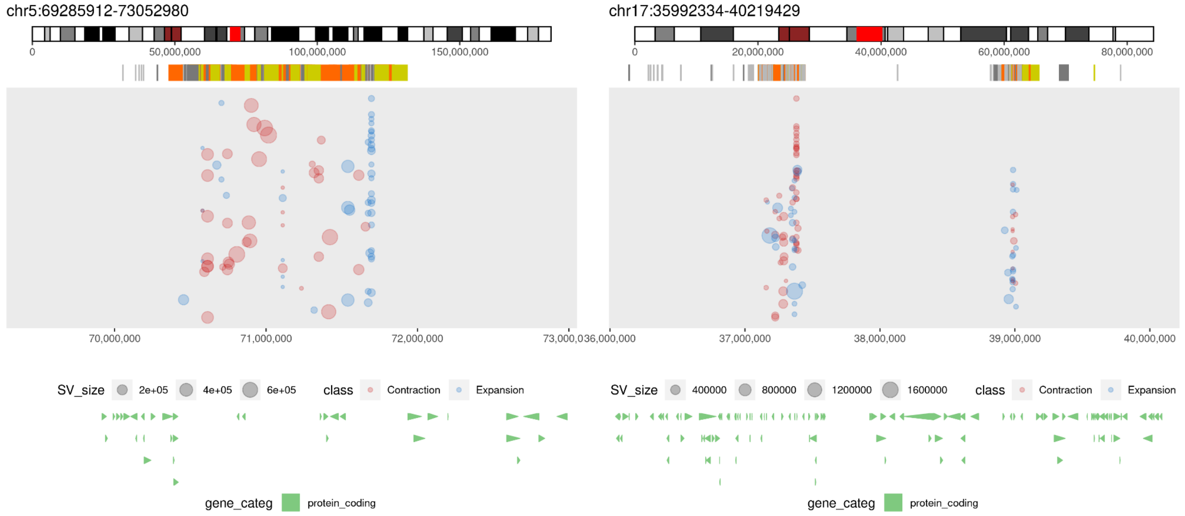


### Figure S18: Example regions with frequent contig alignment discontinuities.

Example regions (left - *SMN1/SMN2* loci; right - *TBC1D3* loci) with frequent expansions and contractions. Each region is highlighted as a red rectangle on the chromosome-specific ideogram (top track). Below there is an SD annotation for a given region represented as a set of rectangles colored by sequence identity. Expansions and contractions of each contig alignment with respect to the reference (T2T-CHM13 v1.1) are depicted as blue and red dots, respectively. The size of each dot represents the size of an event.

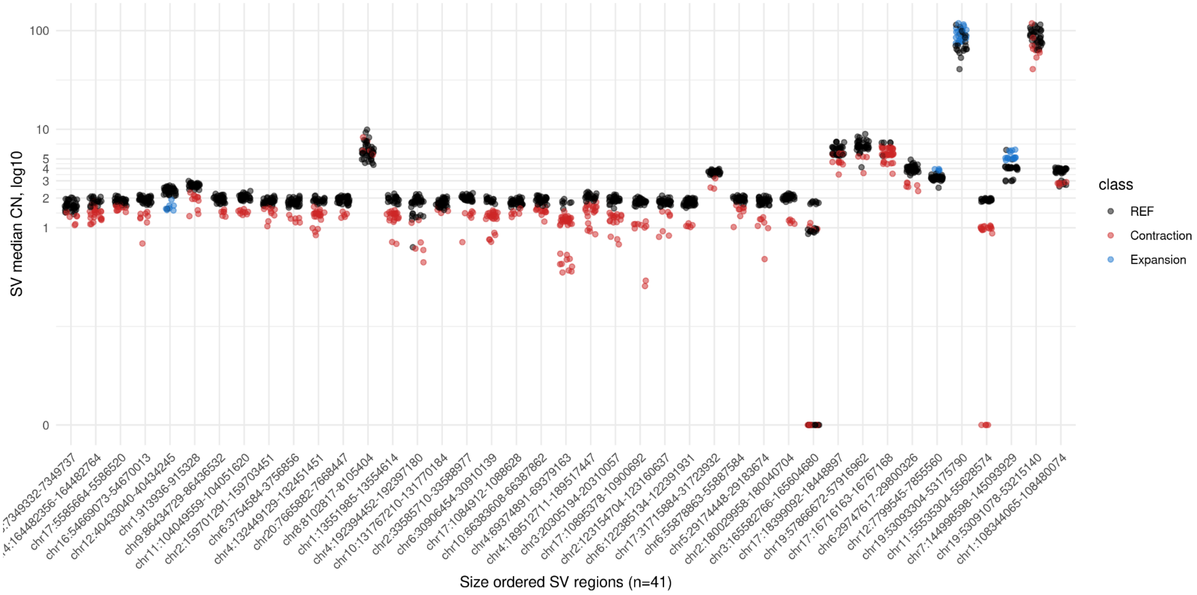


**Figure S19: Contig alignment discontinuities supported by Illumina read-depth.**
A scatterplot of 41 selected regions from frequent contig alignments discontinuities with additional support in short-read copy number (CN) profiles. Each dot represents a short-read median CN in each defined region across all 94 haplotypes. Each dot is colored based on predicted SV class (red - Contraction, blue - Expansion) in a given haplotype. Haplotypes with no predicted SV are colored black (REF - reference).

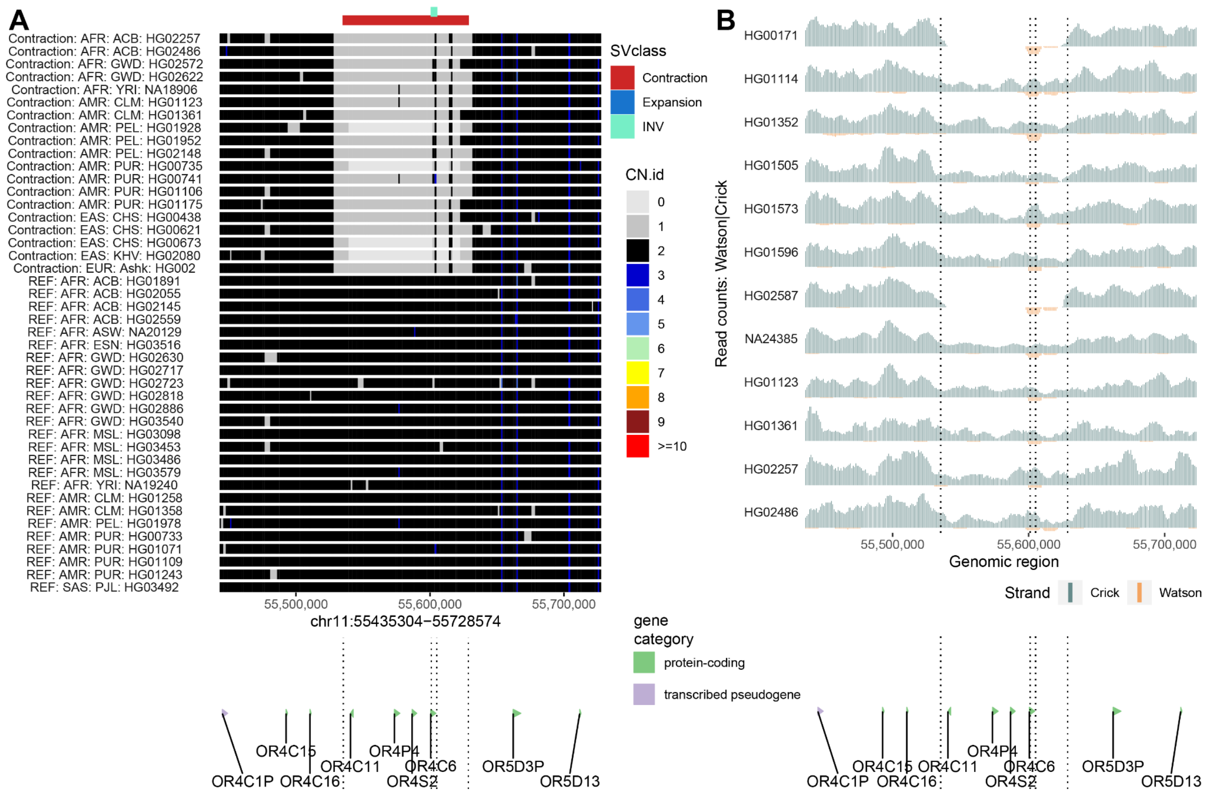

**Figure S20: Inversion-associated microdeletion.**

**A**) Each row represents a short-read CN profile colored by a defined CN. Contracted and inverted regions are highlighted at the top. At the bottom we show protein-coding genes from a given genomic region along with highlighted (dotted lines) breakpoints of contracted and inverted regions (top track). **B**) Read-coverage profiles of Strand-seq data from selected human individuals summarized as binned (bin size: 10 kbp, step size: 1 kbp) read counts represented as bars above (teal; Crick read counts) and below (orange; Watson read counts) midline. Regions with reads aligned to both Watson and Crick reads represent a heterozygous inversion as only one homologue is inverted with respect to the reference. Regions with reads aligned only in Watson orientation represent a homozygous inversion as both homologs are inverted with respect to the reference while regions with purely Crick reads are represented by both homologs being in direct orientation with respect to the reference. Vertical dotted lines highlight the deleted and inverted regions.

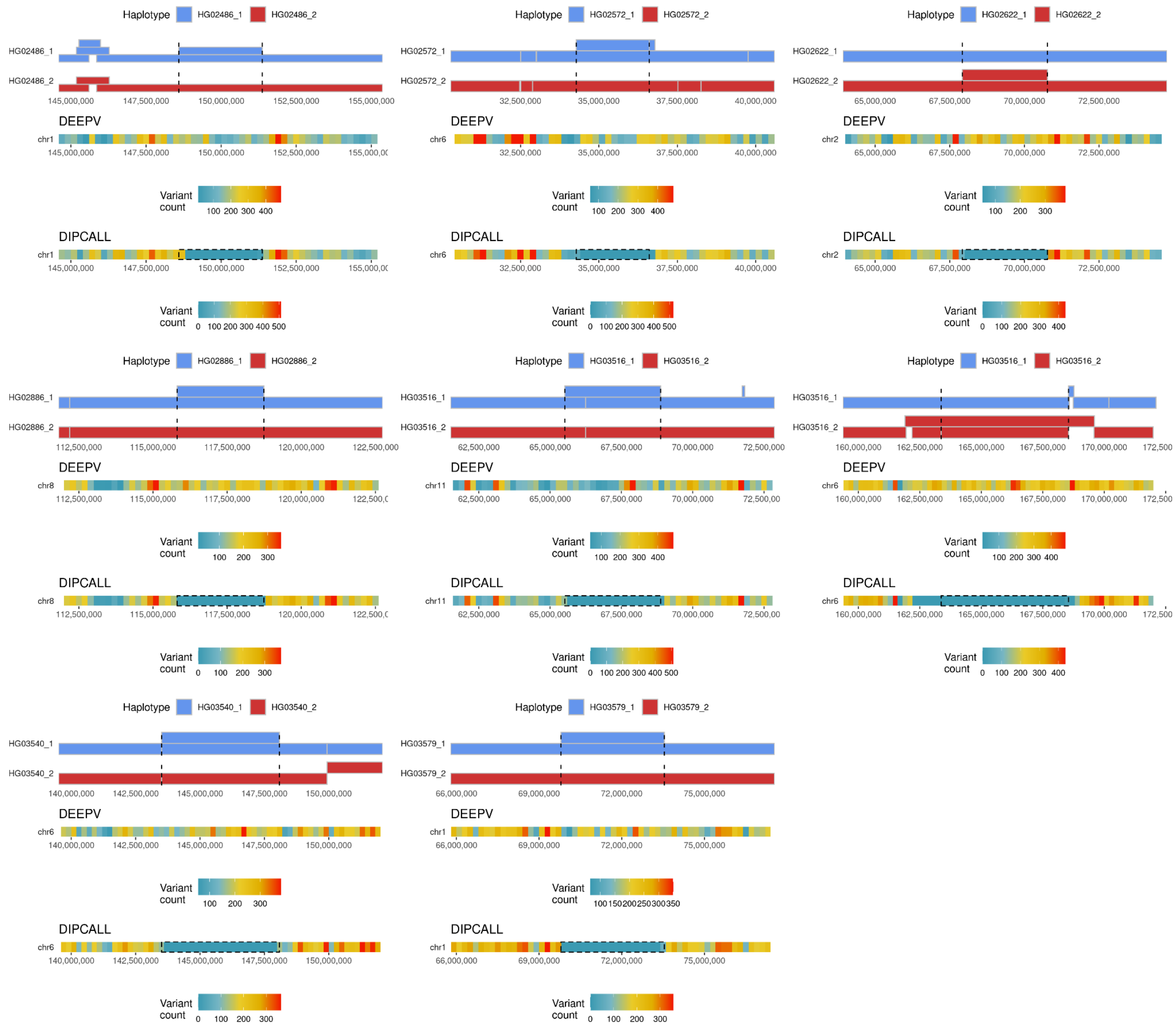


**Figure S21: Biases in SNV discovery over regions with embedded contigs.**
Visualization of large (≥2 Mbp), embedded contigs in a single haplotype and their effect on SNV discovery. Each plot presents data in multiple horizontal tracks. The first track shows contig alignments of a defined region separately for haplotype 1 (blue, paternal) and haplotype 2 (red, maternal). Overlapping contig alignments are stacked on top of each other. The second track shows counts of heterozygous SNVs (in 200 kbp bins) discovered in aligned HiFi reads to the T2T-CHM13 (v1.1) reference (using deepvariant, DEEPV) over a defined region. The third track shows counts of heterozygous SNVs (in 200 kbp bins) discovered in aligned contigs to the GRCh38 reference (using dipcall, DIPCALL) over a defined region. Note: dipcall heterozygous SNVs were lifted from GRCh38 coordinates to T2T-CHM13 (v1.1) coordinates using liftOver (Hinrichs 2006) (**Methods**, **Data availability**).

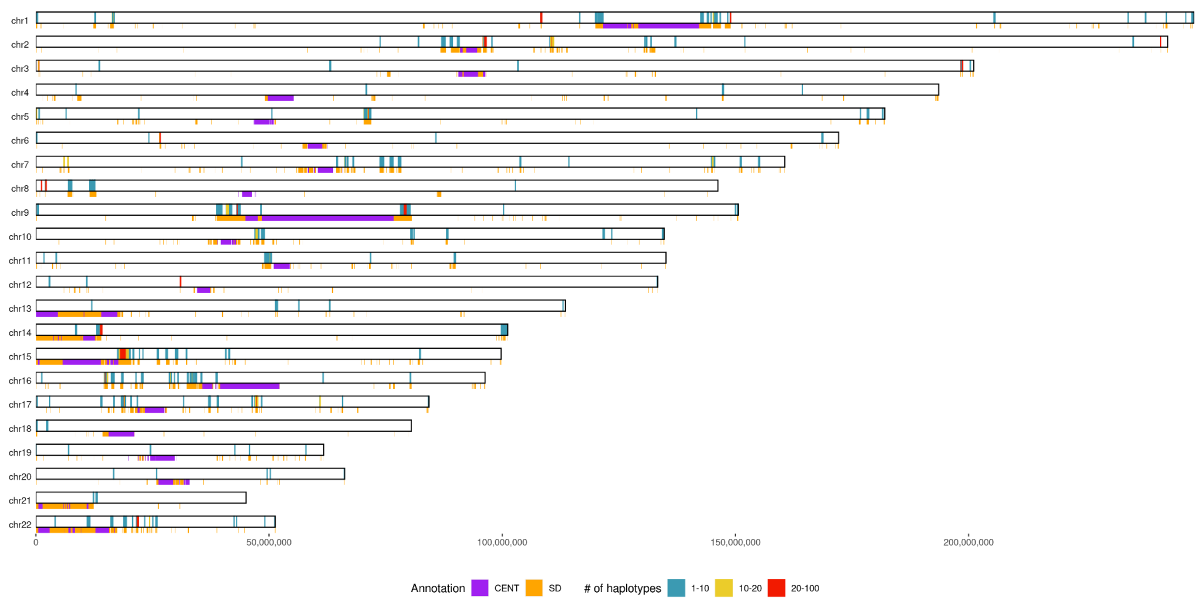


**Figure S22: Genome-wide distribution of detected CNV regions.**

Genome-wide distribution of CNV regions for a subset of 44 HPRC samples (88 haplotypes). Color range reflects the number of haplotypes overlapping each other in any given genomic region (n=255). At the bottom of each chromosomal bar there is annotation of SDs (orange) and higher order centromeric regions (CENSAT, purple).

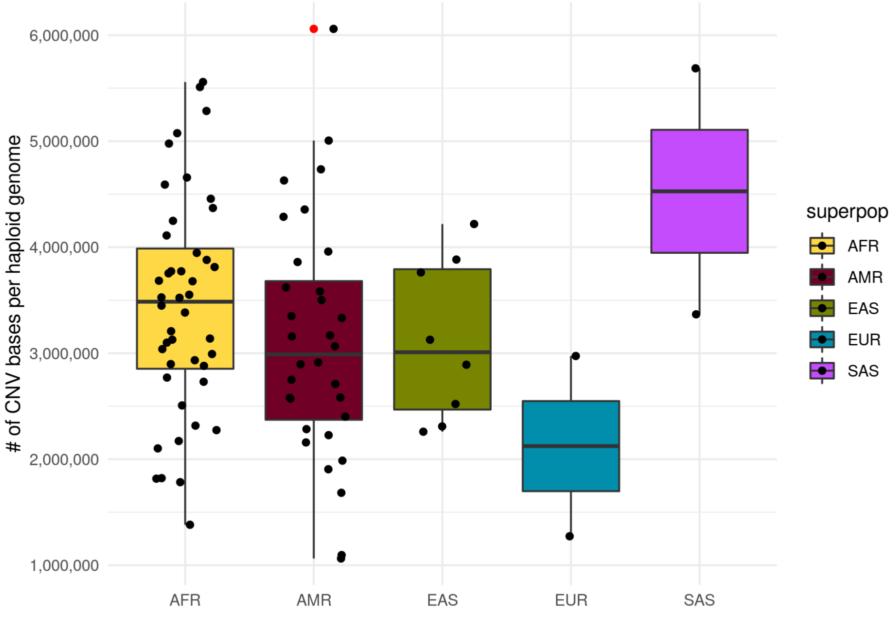


**Figure S23: Genome-wide distribution of detected CNV regions.**

A boxplot showing the distribution CNV bases (extra CN ≤10) per haploid genome and per superpopulation identifier (AFR - African, AMR - American, EAS - East Asian, SAS - South-East Asian and EUR - European).

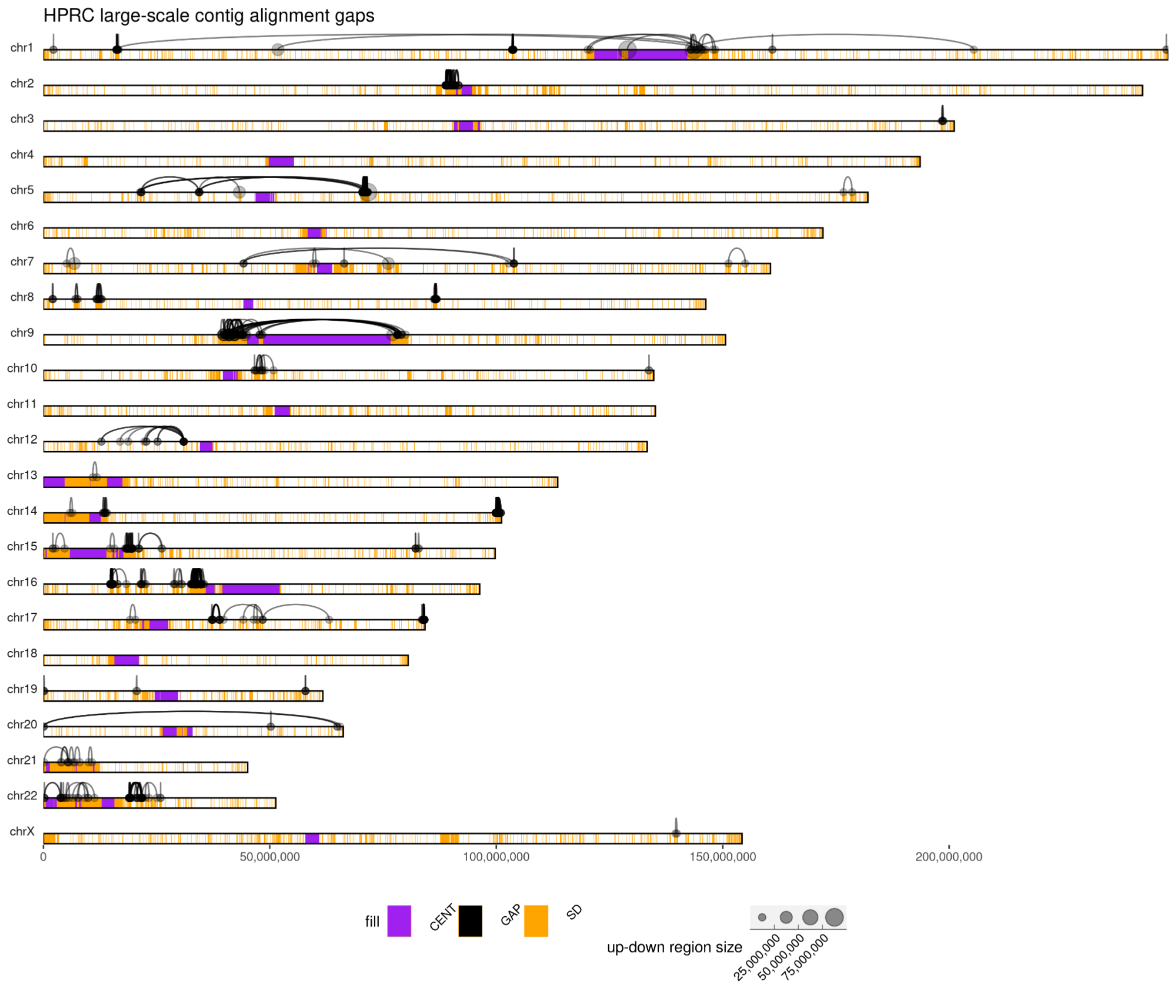


**Figure S24: Genome-wide distribution of split contig alignment ends.**

Each chromosome is depicted as a horizontal bar with the locations of SDs and centromeric regions highlighted as orange and purple rectangles. Contigs divided into multiple contig alignment ends are visualized as links between subsequent pieces of a single contig aligned to the reference (T2T-CHM13 v1.1). The length of the aligned piece of a contig is defined by the size of each dot while the width of each link depicts the size of a gap between aligned pieces of a given contig.

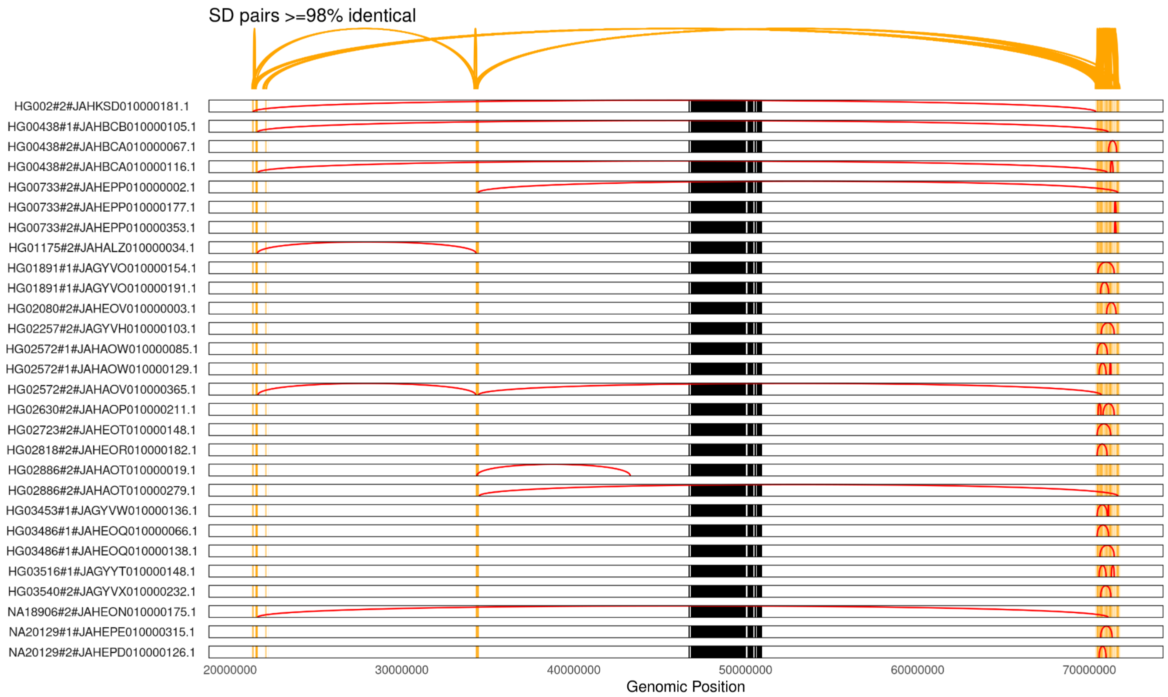


### Figure S25: Split contig alignment ends mapping to SMN gene region.

The SMN gene region on chromosome 5 is visible at the far right end of the plot. The chromosome 5 centromere is in the middle highlighted as black rectangle. Each contig assembly with contig alignment discontinuity larger than 1 Mbp in the SMN gene region is plotted per row. Red lines are representing gaps connecting two pieces of a single contig that maps far apart with respect to the reference (T2T-CHM13 v1.1). The complexity of this region is also visible at the genome-wide plot above.

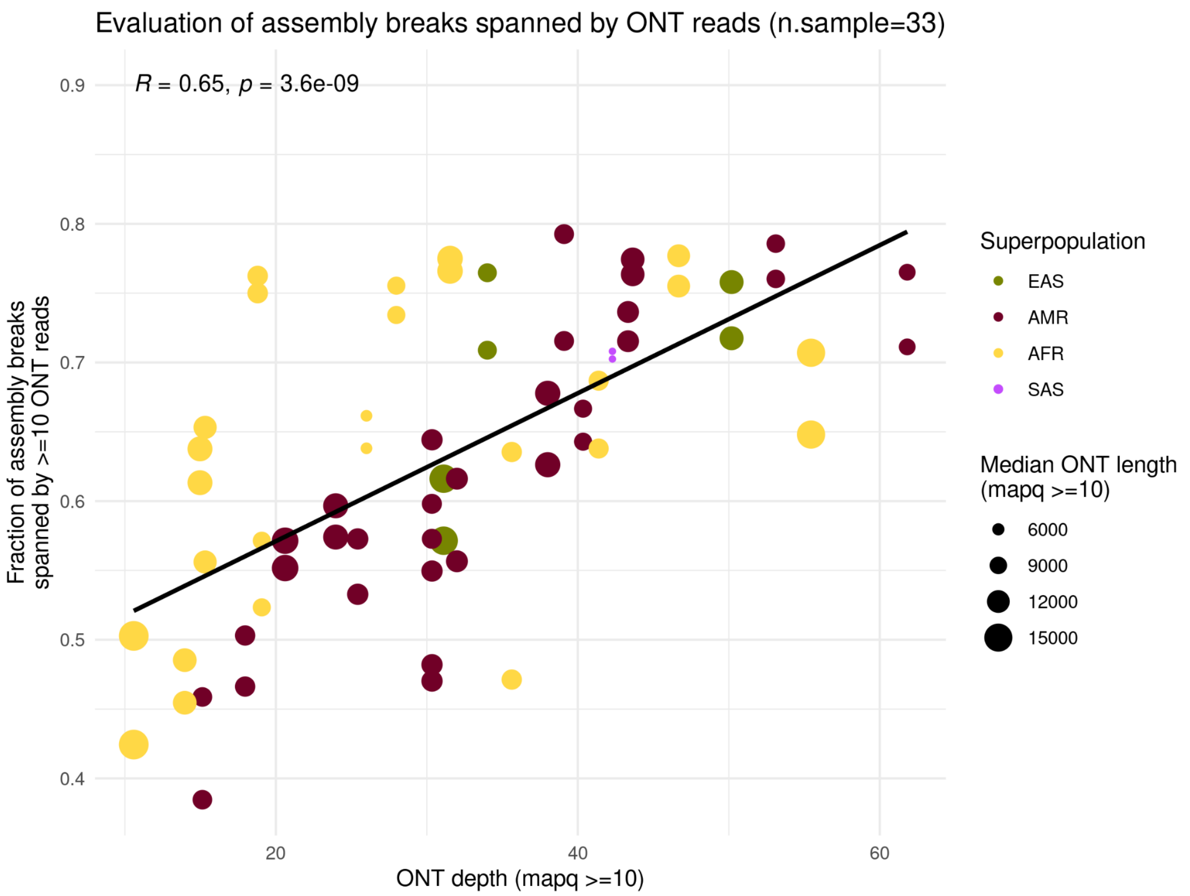


**Figure S26: Extent of assembly gap regions spanned by ONT reads.**

Each dot represents a sample (n=33) specific haplotype (n=66) with a fraction of assembly gap regions spanned by ONT reads on the y-axis and ONT read depth on the x-axis. The size of each dot reflects the median length of ONT reads. Each dot is colored by the superpopulation any given sample originates from. Black line highlights fitted linear model.

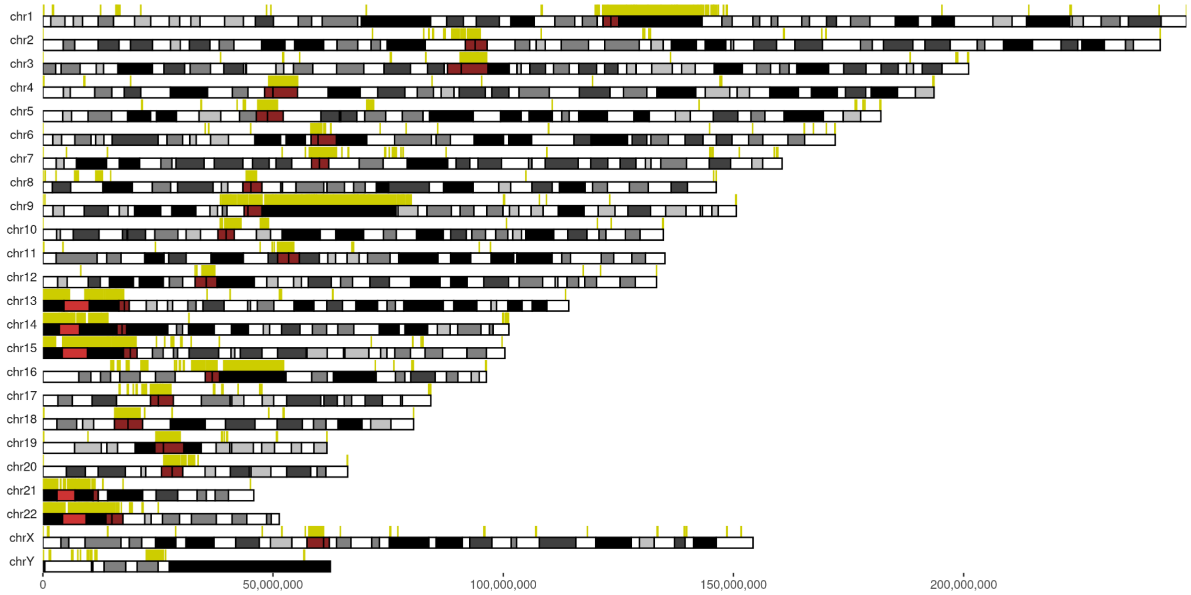


**Figure S27: Genome-wide distribution of defined 'brnn' sites.**
Defined brnn sites (n=638) are visualized as yellow rectangles above each chromosomal ideogram.

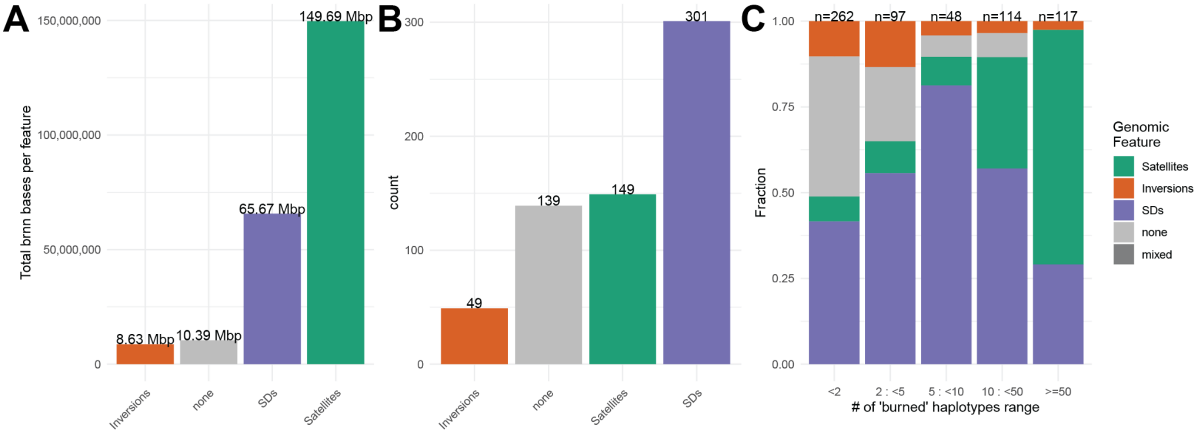


**Figure S28: Genomic feature proportions per 'brnn' site.**
**A**) The total number of base pairs of various genomic features (Inversions, SDs, Satellites, and 'none' of those) overlapping a nonredundant set of 'brnn' regions (n=638). **B**) An assignment of each 'brnn' region based on a single majority genomic feature (proportion of the given feature >0.5). **C**) Proportion of genomic features assigned to each brnn region based on the number of 'burned' haplotypes within each brnn region.

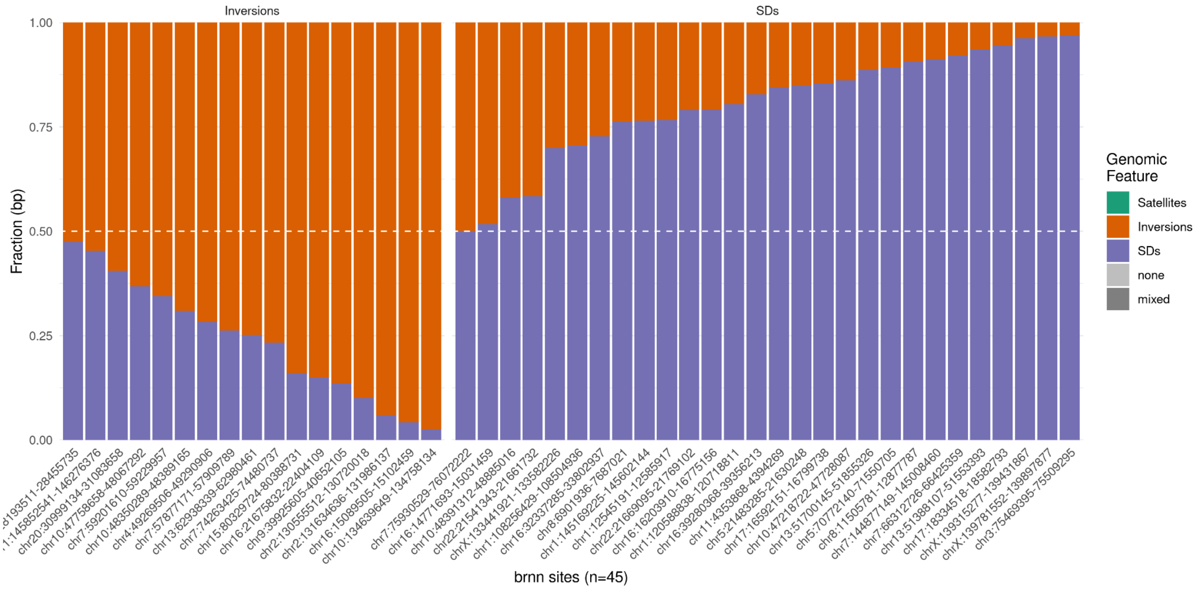


### Figure S29: Brnn sites at the boundaries of SD flanked inversions.

The proportion of base pairs colored by tested genomic features ('none' - zero overlap with tested features) that overlaps brnn regions reported at the boundaries of SD-flanked inversions (n=45). Regions are stratified and ordered by the majority genomic feature.

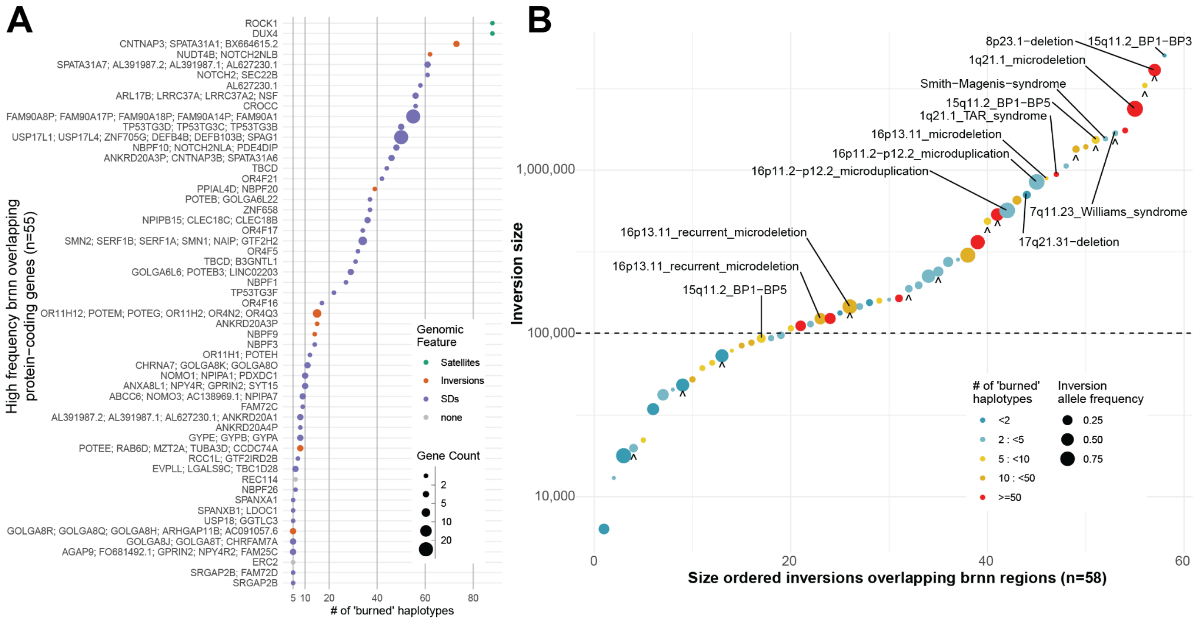


### Figure S30: Importance of regions excluded from the pangenome graph.

**A**) Examples of protein-coding genes overlapping frequent brnn regions (≥5 'burned' haplotypes, n=279) colored by the assigned genomic feature. **B**) Size distribution (ordered x-axis), allele frequency (point size), and brnn frequency (point color) of all inversion sites (n=58) with a detected overlap (10%) with the 'brnn' regions. Inversion previously marked as recurrent are marked by **'^'** sign (Porubsky et al. 2022).

### SUPPLEMENTAL TABLES (1-9):

**Table S1: Regions of wrongly resolved homozygous inversions. [.xlsx file]**
**Table S2: List of all defined simple contig ends. [.xlsx file]**
**Table S3: Protein-coding genes overlapping frequent assembly gaps in HPRC assemblies. [.xlsx file]**
**Table S4: List of frequent assembly gaps in HPRC assemblies. [.xlsx file]**
**Table S5: List of very high frequency assembly gaps in HPRC assemblies. [.xlsx file]**
**Table S6: List of high frequency contraction and expansions. [.xlsx file]**
**Table S7: List of CNV regions at multiple contig mappings. [.xlsx file]**
**Table S8: List of inversions overlapping brnn regions. [.xlsx file]**
**Table S9: Locations of ONT data. [.xlsx file]**

### SUPPLEMENTAL NOTES

There are regions in current genome assemblies that are either completely missing or incorrectly assembled which pose a problem for construction of pangenome graphs. We projected such regions on T2T-CHM13 (v1.1) reference and identified 638 nonoverlapping regions that have been excluded from the pangenome graph construction at least once and up to 88 times (**Fig. S27, Methods**). As expected, the majority of excluded bases corresponds to satellite DNA (~149.7 Mbp) and highly identical SDs (~65.7 Mbp) (**Fig. S28A**). Based on the majority overlap, we observe an opposite trend with SD bases underlying most (n=301) of the ‘brnn’ regions followed by satellite DNA (n=149) (**Fig. S28B**). In line with sequence properties of satellite DNA and SDs, we found SD regions being excluded at a lower frequency (>5 : <50) while satellite DNA is excluded from the pangenome graph most frequently (>50) (**Fig. 28C**). When considering only frequently burned regions (≥5, n=279), we found 55 ‘brnn’ regions that overlap a total of 171 protein-coding genes, the majority of which (n=43) lie within SD-rich regions (**Fig. 30A**). As outlined above, we again observe inversions being associated with regions that complicate pangenome graph construction (~8.6 Mbp, n=49) (**Fig. 28A-B**). This is because large polymorphic inversions are often flanked by highly identical SDs supported by a number of ‘brnn’ regions overlapping both inversions and SDs (**Fig. S29**). There are 58 inversion sites (median size of 160 kbp) overlapping ‘brnn’ regions of which 14 are associated with morbid CNVs and 13 have been labeled as recurrent in the recent study (**Fig. S30B, Table S8**).
